# Artificial intelligence-augmented drug discovery identifies gefitinib as a potential treatment for ALS

**DOI:** 10.1101/2025.03.06.641147

**Authors:** Monika A. Myszczynska, Matthew J. Stopford, Nóra M. Márkus, Sophie E. L. Nyberg, Nicole L. Stone, Sarah M. Granger, Allan C. Shaw, Raquel Rua Martins, Chloe F. Allen, Amy F.A. Keerie, Tyler R. Wells, Ruth H.E. Thomas, Sian H. Brown-Wright, David W. Sheppard, Anne Phelan, Daniel P. Smith, Peter J. Richardson, Richard J. Mead, Laura Ferraiuolo

**Affiliations:** Sheffield Institute for Translational Neuroscience, Department of Neuroscience, University of Sheffield, 385a Glossop Road, Sheffield S10 2HQ, United Kingdom; BenevolentAI, 4–8 Maple Street, London, W1T 5HD, United Kingdom

**Author notes:** Corresponding author: Laura Ferraiuolo, **Email:**. These authors contributed equally (Monika A. Myszczynska, Matthew J. Stopford). These authors contributed equally (Richard J. Mead, Laura Ferraiuolo). **Author Contributions:** MAM, MJS, NMM, DWS, AP, PJR, RJM, LF designed research; MAM, MJS, NMM, SELN, NLS, SMG, ACS, RRM, CFA, AFAK, TRW, RHET, SHBW performed research; DWS, AP, DPS, PJR contributed new reagents/analytic tools; MAM, MJS, NMM, SELN, NLS, SMG, RRM, RJM, LF analysed data; MAM, MJS, NMM, DPS, PJR, LF wrote the paper; PJR, RJM, LF edited the paper with contribution from all authors. **Competing Interest Statement:** The research described in this manuscript was funded by BenevolentAI. DWS, AP, DPS and PJR were employed by BenevolentAI. **Classification:** Biological sciences; Neuroscience.

**Keywords:** amyotrophic lateral sclerosis (ALS), artificial intelligence (AI), astrocytes, TDP-43, autophagy

## Abstract

Amyotrophic lateral sclerosis (ALS) is characterised by motor neuron (MN) death; however, astrocytes play a key role in disease pathogenesis. Developments in the field of artificial intelligence (AI) have the potential to impact drug discovery in multiple ways, including the rapid identification of drug repurposing candidates. A combination of natural language processing and deep learning algorithms was used to generate a knowledge graph based on scientific literature, omics and chemical databases, and other public sources with the aim to identify drug repurposing candidates for ALS. The aim of the study was to determine the effect of a cancer compound identified by AI, gefitinib, on MN survival, and to decipher its mode of action in *in vitro* and *in vivo* models of ALS. We used co-cultures of healthy motor neurons with ALS patient-derived astrocytes (iAstrocytes), obtained through a semi-direct conversion protocol, to assess the neuroprotective properties of gefitinib. Compound treatment led to a significant rescue of MNs cultured with ALS iAstrocytes and a significant reduction in the levels of cleaved TDP-43 fragments in ALS iAstrocytes. Our data suggest that gefitinib-mediated activation of autophagy decreased the 35 kDa fragments of TDP-43. In a proof-of-concept *in vivo* study in SOD1^G93A^ mice, gefitinib treatment significantly delayed the onset of neurological symptoms, thus showing the potential of AI-augmented drug discovery for neurodegenerative disorders.

**Significance Statement:** This study presents an AI-augmented method of identifying potential repurposing candidates for disease with an unprecedented speed. The AI’s results were validated *in vitro* using iAstrocytes differentiated from induced neuronal progenitor cells (iNPCs), which are pathophysiologically relevant models suitable for studying neurodegeneration. iNPCs recapitulate many pathological hallmarks of the disease and they retain the ageing phenotype of the patient that they are obtained from. TDP-43 proteinopathy is one of the disease hallmarks observed in patients and is present in 97% of ALS patients. Here, we show gefitinib, a repurposing candidate identified by AI, improves survival of MNs in a co-culture with patient-derived astrocytes and can modulate TDP-43 proteinopathy.

## Introduction

Amyotrophic lateral sclerosis (ALS) is a fatal neurodegenerative disease characterised by loss of upper and lower motor neurons (MNs), progressive muscle weakness and wasting. The median survival after diagnosis is 2–3 years, with respiratory failure being the most common cause of death (1). Riluzole and edaravone are the only Food and Drug Administration (FDA)-approved drugs for sporadic ALS. These drugs show marginal benefits in terms of improved survival or slowing of decline in disease progression. Therefore, identifying new drugs for ALS remains an urgent unmet need.

Approximately 90% of ALS patients have no family history of the disease and are classed as sporadic (sALS). sALS remains critically understudied compared with familial ALS (fALS) due to lack of good models. sALS and fALS are clinically indistinguishable, which suggests a presence of overlapping pathological mechanisms. Currently, a GGGGCC (G_4_C_2_) hexanucleotide repeat expansion in the first intron of the C9orf72 gene (2) is the most common genetic cause of ALS; the mutation is observed in both fALS and sALS cases (3), and accounts for 45% of fALS in western European populations, 4% in Black African and 8% in Hispanic populations and 5–10% of sALS cases (4–6). Physiologically, healthy individuals carry up to 23 copies of the G_4_C_2_ repeat – possessing more than 23 copies is pathogenic (7). However, no clear relationship exists between the length of the expansion and the severity of the disease (8).

While the full spectrum of its physiological functions is still being investigated, the role of C9orf72 protein is linked to initiation of autophagy (9), vesicle formation and trafficking (10).

ALS is a multimechanistic disorder. The pathogenic process and mechanism of disease onset and progression remain poorly understood, though dysregulation of RNA processing and metabolism, glutamate excitotoxicity, oxidative stress, axonal transport impairment, and mitochondrial dysfunction are some of the well-described common factors (11, 12).

In particular, transactive response DNA binding protein 43 (TDP-43) inclusions are a hallmark of ALS pathology and are observed in 97% of cases; notably, TDP-43 inclusions are not observed in patients with *SOD1* or *FUS* mutations (13, 14). TDP-43 is located mostly in the nucleus in healthy individuals where it is involved in RNA metabolism and processing (15). In times of cellular stress, TDP-43 is incorporated into cytoplasmic stress granules, temporary inclusions that sequester the most essential mRNA molecules for translation in order for the cell to survive the stress event (16,17). However, TDP-43 may remain in the cytoplasm and form aggregates due to nuclear export/loss, lack of nuclear import, protein cleavage and phosphorylation of the C-terminal domain (18–20). Cytoplasmic TDP-43 in ALS patients is truncated into fragments of 35, and 18–25 kDa, all containing its C-terminal domain, which are often phosphorylated, ubiquitinated, or both (21). The precise mechanism triggering the mislocalisation and aggregation of TDP-43 remains unknown, similarly to the contribution of these events to MN death (22, 23). Because of its widespread presence in ALS cases, TDP-43 proteinopathy is the target and readout of several drug discovery efforts (24–27).

Developing new drugs is costly and time-consuming (28). Drug repurposing applies existing drugs with well understood safety and efficacy profiles to new diseases, which can significantly shorten drug development times and makes it an attractive alternative to novel molecule discovery (29). To further improve and speed up this process, machine learning approaches have been investigated as potential clinical aids (30) as well as drug repurposing (31) and drug discovery tools (32). Recent efforts focused on identifying new therapeutic targets and designing *de novo* compounds that engage with a druggable pocket of the target (33), retrosynthesis and reaction prediction (34), and virtual screenings (35). These methods have so far been used for target identification, inhibitor discovery, and multitarget drug discovery in Parkinson’s disease (36–39), Alzheimer’s disease (40–43), and neuropsychiatric disorders (44). Machine learning can benefit drug repurposing by augmenting the speed with which a known drug is assigned to a new disease environment by matching the drug’s targets to genetic signatures of the disease. Although still in its infancy, such methods identified baricitinib, an alopecia and rheumatoid arthritis drug, as a treatment for COVID-19, demonstrating their utility (45).

In this study, we identified five compounds with a therapeutic potential for ALS using artificial intelligence (AI). Of the five candidates, one compound, gefitinib, rescued MN survival in co-culture with sporadic, SOD1 and C9orf72-ALS patient-derived astrocytes and reduced the levels of TDP-43 fragments *in vitro* via autophagy activation. Enhanced central nervous system (CNS) exposure of gefitinib (mediated by PGP inhibitor elacridar) in SOD1^G93A^ mice delayed the neurological symptom onset and was associated with a trend towards a decrease in misfolded SOD1 accumulation.

## Results

### Artificial Intelligence technology augments ALS therapeutic discovery

We used proprietary AI technology to augment drug discovery in ALS. AI was employed to process a vast corpus of scientific literature and synthesise it with existing structured biomedical data in the BenevolentAI knowledge graph (46, 47). A combination of natural language processing algorithms and deep learning algorithms were used to read and process scientific literature and patents (Fig. 1A). As a result, “mined” unstructured information was identified and added into a knowledge graph, to sit alongside the existing structured and curated information available from public biomedical databases. This data handling process semantically integrated structured and unstructured biomedical data, and the resulting knowledge graph was used to generate hypotheses for ALS therapeutics.

**Figure 1.**
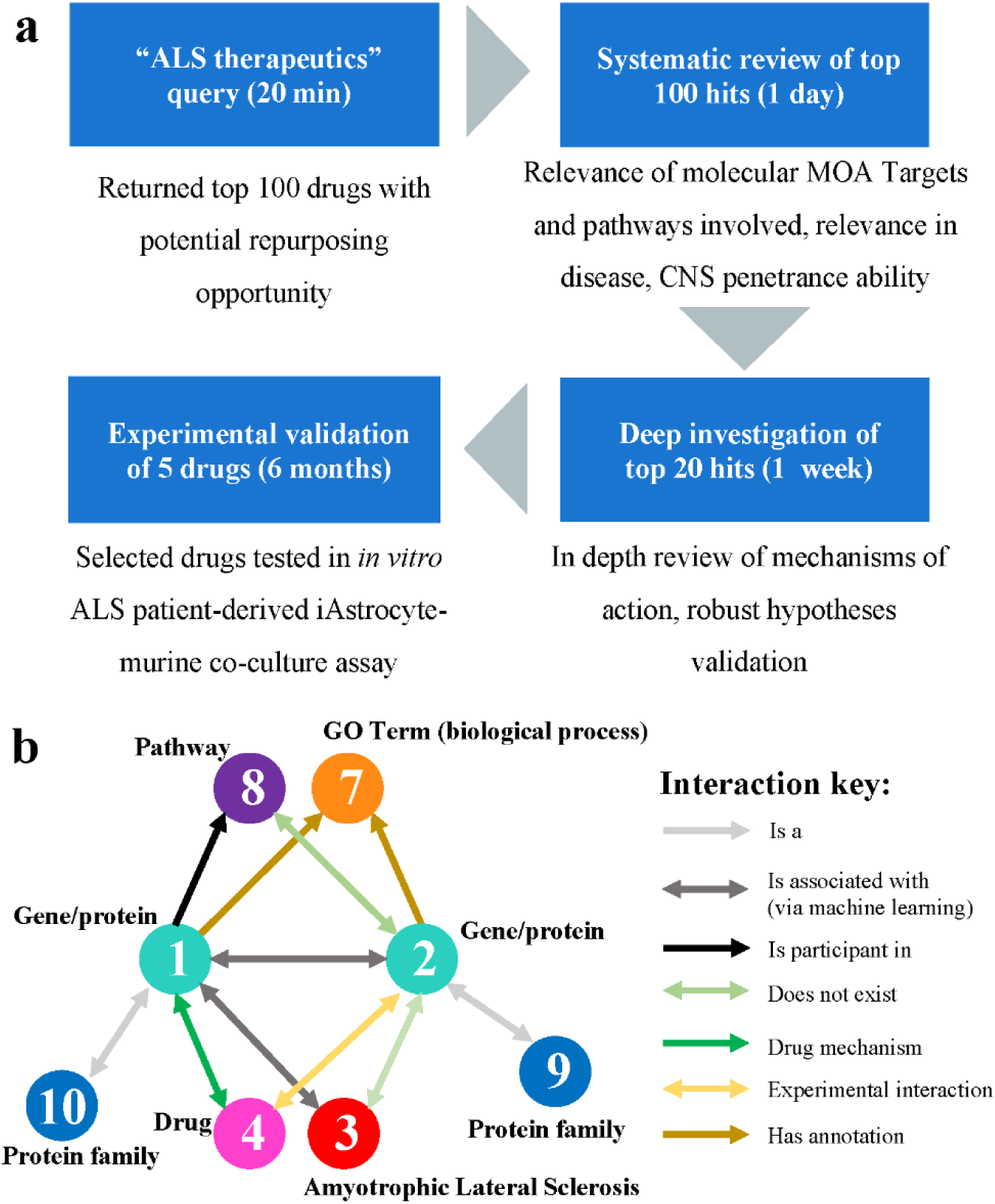
Artificial Intelligence augmented ALS drug discovery. **a)** An ALS therapeutics query was run on BenevolentAI’s knowledge graph, which identifies potential ALS therapeutic drugs by running the following logic: Given Amyotrophic Lateral Sclerosis (3); Find a set of Generics (gene/proteins) (1) that have been found to be associated to Amyotrophic Lateral Sclerosis (3) via our AI. Find a second set of Generics (gene/proteins) (2) that have been associated with the first set of Generics via our AI. Make sure this second set of Generics (2), have *never been associated* with Amyotrophic Lateral Sclerosis (3) across any data source or our AI. Make sure that both sets of Generics (1 & 2) belong to different Protein Families (9 & 10). Make sure that both sets of Generics (1 & 2) have been annotated as taking part in the same biological process (7). Make sure that the second set of Generics (2) do not appear on the same biological pathways (8), as the first set of Generics (2). Make sure that one set of Generics (1) are the primary mechanism of a drug (4). Make sure that the other set of Generics (2) have a recorded experimental interaction with the same drug (4). **b)** The ALS therapeutics query identified 100 drugs for potential repurposing for ALS. These hits were systematically reviewed and then validated using biological models of ALS.

Briefly, relation extraction from unstructured data focused on linkages such as Encode-Attend-Tag relationships (EAT), a proprietary term used to describe relationships that demonstrate biological association between two biomedical concepts, in this case diseases and proteins. EAT relationships were extracted by an algorithm that uses distant supervision to learn syntactical association on a biomedical literature database in order to extract relationships that provide evidence for a particular Disease-Protein or Gene link. In addition to identifying supporting sentences in the literature, this process assigns a confidence score to each relationship that reflects the accuracy of that Disease-Protein link. Other relationships included in the knowledge graph include Literature-based Therapeutic Evidence edges that convey a therapeutic relationship and Syntactic Subject-Verb-Object relationships that capture directional biological associations (46).

The resulting AI-enriched knowledge graph was then queried using an interactive knowledge graph pattern creation tool, which provides flexibility to ask a situational question of ALS that comprised the disease, biological processes, any number of proteins and relationships and can exclude specific relationships. The ALS query required a human to sketch a knowledge graph pattern comprising entities of interest, such as diseases, processes, pathways, genes, proteins, and drugs. Only one part of the pattern was specified (Disease = ALS) and all other parts of the pattern were left as “unknowns” for the search algorithm to complete. Using this tool, we searched for pairs of interacting proteins only one of which was associated with ALS. We specified that the proteins must also be associated with the same biological processes and pathways, and then looked for drugs which modulated the activity of both proteins (Fig. 1B). We ran this ALS therapeutics knowledge pattern, as well as four other variations of this pattern that brought in cell-specific processes to add biological context, and pathway analysis to find selective targets. Importantly, the ALS therapeutics knowledge patterns were designed to identify drugs that can act on multiple dysfunctional pathways in ALS. The combined results were in the form of an aggregated spreadsheet, where the frequency of appearances of the drugs and targets were tallied. This process identified 100 compounds with potential repurposing opportunity and took 20 minutes for the algorithm to triage. These 100 hits were systematically reviewed by subject matter experts (SMEs), and hits were ranked with regard to mechanism of action (MOA) relevance, the targets and pathways involved, the relevance in disease, and potential for CNS penetrance An in-depth review of MOA and robust hypothesis validation were performed, which entailed cross validating the compounds on new knowledge graphs, using the top 20 hits based on the ranking, and after one week, five compounds (ambroxol, erlotinib, gefitinib, RTA408, and siramesine) were selected for experimental validation. Gefitinib and erlotinib were predicted to alter multiple signalling pathways including those involved in neuroinflammation and autophagy, ambroxol to reduce MN excitability, RTA408 to induce antioxidant defences and siramesine to affect endoplasmic reticulum stress pathways.

### Gefitinib improves MN survival in co-culture with C9orf72-ALS, sALS, and SOD1-ALS iAstrocytes

To validate their therapeutic potential, ambroxol, erlotinib, gefitinib, RTA408, and siramesine were tested in a pathophysiologically relevant *in vitro* co-culture model of ALS, which recapitulates the non-cell autonomous mechanism of endogenous toxicity of astrocytes in the diseases (48). This rapid, highly reproducible methodology uses induced neural progenitor cells (iNPCs), reprogrammed from ALS patient and control fibroblasts (49–53). Here, we differentiated three C9orf72-ALS patient (C9_1, C9_2, C9_3) and three healthy control (Con_1, Con_2, Con_3) iNPC lines into induced astrocytes (iAstrocytes) and co-cultured them with naïve mouse embryonic HB9-GFP+ MNs (54, Fig. 2A). When co-cultured with C9orf72-ALS patient iAstrocytes, MN survival was significantly reduced compared to a co-culture with control iAstrocytes (56.8–79.4% reduction depending on the patient iAstrocyte line, P = 0.001; Fig. 2B).

**Figure 2.**
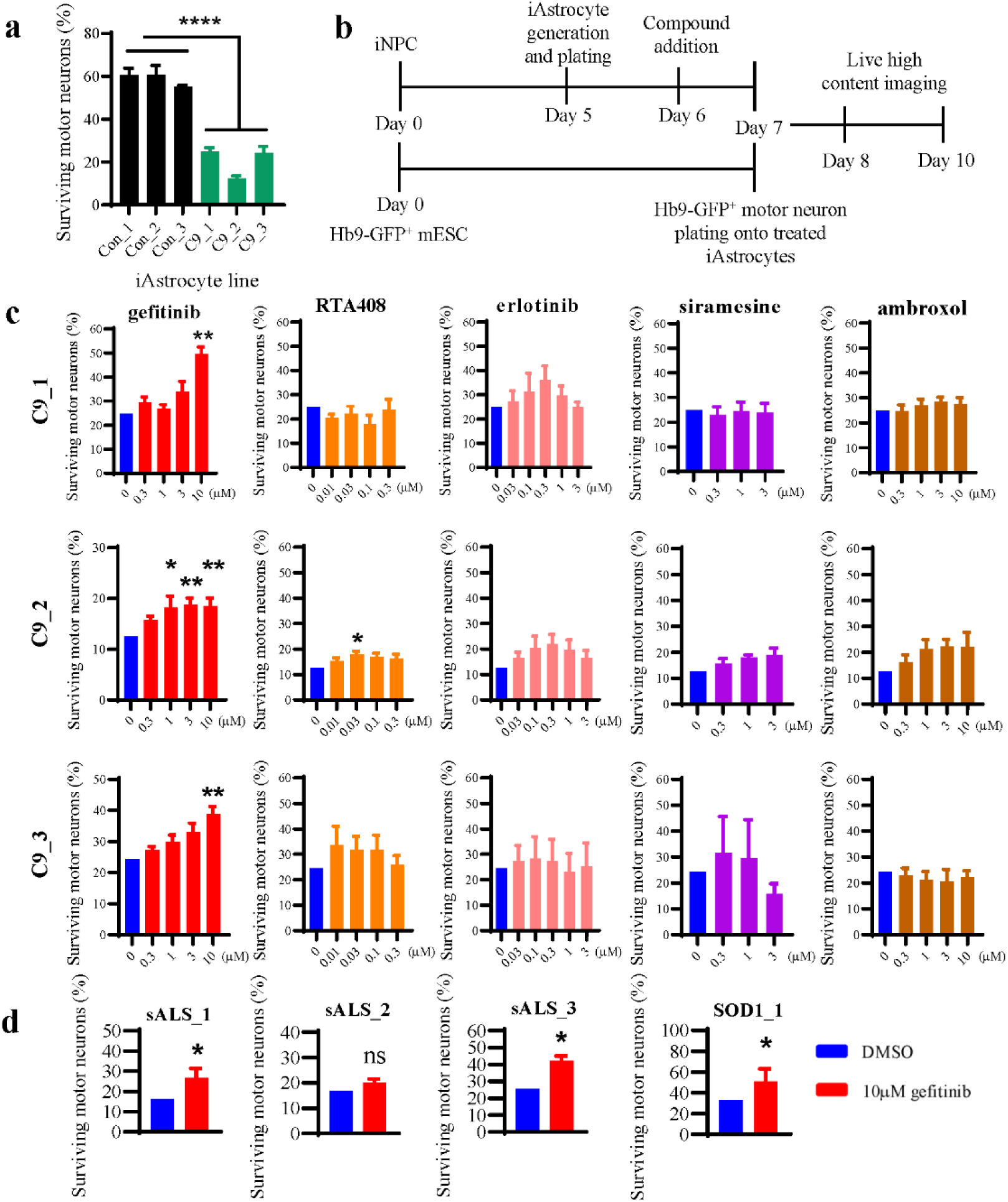
Gefitinib improves motor neuron survival in co-culture with ALS patient-derived iAstrocytes. **a)** Control and C9orf72-ALS patient-derived iAstrocytes were co-cultured with murine Hb9-GFP^+^ MN. % MN survival was calculated as number of MN at 96 h as a percentage of number of MN at 24 h. Data = mean ± SEM; n = 7-13 (except Con_3 n=2, excluded from statistical analysis); One-way ANOVA with Tukey’s multiple comparisons test, ****P<0.0001. **b)** Schematic of the high-throughput human iAstrocyte-murine Hb9-GFP^+^ MN co-culture assay. **c)** C9orf72-ALS patient-derived iAstrocytes were treated with a range of gefitinib doses, and co-cultured with murine Hb9-GFP^+^ MN. % MN survival was calculated as number of MN at 96 h as a percentage of number of MN at 24 h, and then normalised to the DMSO control value. Data = mean ± SEM; n = 3-7; Kruskal-Wallis test with Dunn’s multiple comparison’s test; *P < 0.05; **P < 0.01; ***P < 0.001. **d)** sALS and SOD1-ALS patient-derived iAstrocytes were treated with 10 µM gefitinib, and co-cultured with murine Hb9-GFP^+^ MN. % MN survival was calculated as number of MN at 96 h as a percentage of number of MN at 24 h, and then normalised to the average of DMSO control value. Data = mean ± SEM, n=4-6. Data normalised to DMSO = 100. Mann-Whitney test, *P < 0.05.

Next, we introduced the five compounds into the co-culture system as described before (Fig. 2A; 54). Here, we observed improved MN survival in co-culture with gefitinib-treated C9orf72-ALS iAstrocytes lines in a dose-depended manner, with 10 μM concentration performing best across all lines (Fig. 2C). RTA408 did not improve MN survival at all in co-culture with C9_1 and C9_3 iAstrocytes, however, 0.03 µM RTA408 did improve MN survival by 42.5 ± 23.5% (P = 0.0312) in co-culture with C9_2. Ambroxol, erlotinib, and siramesine did not significantly improve MN survival in co-culture with any of the three C9orf72-ALS iAstrocyte lines (Fig. 2C). Therefore, gefitinib was taken forward for further investigation, as it was the only compound to improve MN survival across all C9orf72-ALS iAstrocyte lines tested. Based on the concentration response among these lines, 10 µM gefitinib was selected for all further investigations and thereafter all administered gefitinib treatments use this concentration for 48 h.

To validate if gefitinib neuroprotective effects could be applied more broadly to sporadic and SOD1-ALS patients, three sALS and one SOD1-ALS iAstrocyte lines were used in the co-culture workflow described above. With the exception of one sALS line, i.e. sALS_2, iAstrocytes, treatment with 10 µM gefitinib led to a significant rescue of MN survival in co-cultures with sALS_1 (P = 0.048), sALS_3 (P = 0.029), and SOD1_1 (P = 0.029) iAstrocytes (Fig. 2D).

Since gefitinib is primarily used for treatment of non-small cell lung cancer (NSCLC) as an inhibitor of the epidermal growth factor receptor (EGFR), inhibition of which has been previously demonstrated to promote neuroprotection (55), we measured the protein expression levels of total EGFR and EGFR phosphorylated at one of its autophosphorylation sites, Tyr-1068 (pEGFR). Elevated levels of EGFR were observed in donor C9_3, whilst EGFR expression in C9_1 and C9_2 was comparable with the control iAstrocytes (Fig. S1A-B). The baseline level of pEGFR in iAstrocytes was generally low and highly variable (Fig. S1A, C). Treatment of iAstrocytes for 48 h with 10 µM gefitinib showed a trend towards pEGFR inhibition, whilst also showing a trend towards an increase in the total EGFR protein expression (Fig. S1D-F), albeit not statistically significant. To further test the involvement of EGFR in promoting MN survival in a co-culture with C9orf72 iAstrocytes, we knocked-down EGFR using an adenovirus expressing a validated shRNA construct against EGFR. EGFR knockdown was achieved (Fig. S1G-H), but had no effect on MN rescue, suggesting that gefitinib causes neuroprotection via a mechanism not involving its primary target (Fig. S1I).

### Gefitinib reduces TDP-43 fragmentation

Given MN rescue in co-cultures with iAstrocytes treated with gefitinib, we hypothesised that gefitinib targets a common pathogenic ALS mechanism. Since TDP-43 proteinopathy is a hallmark of ALS and is observed in 97% of non-FUS and –SOD1 cases (13, 14), we proceeded to characterise the effect of gefitinib on the endogenous TDP-43 fragmentation in iAstrocytes. TDP-43 fragmentation leads to the production of C-terminal-fragments, including a 35 kDa fragment, which is reliably detectable via western blotting in this model (TDP-35, Fig. 3A and B). In addition, all C9orf72 and sALS iAstrocytes, with the exception of C9_2, showed significantly decreased levels of full length TDP-43 compared to control iAstrocytes (P = 0.005, Fig. 3C).

**Figure 3.**
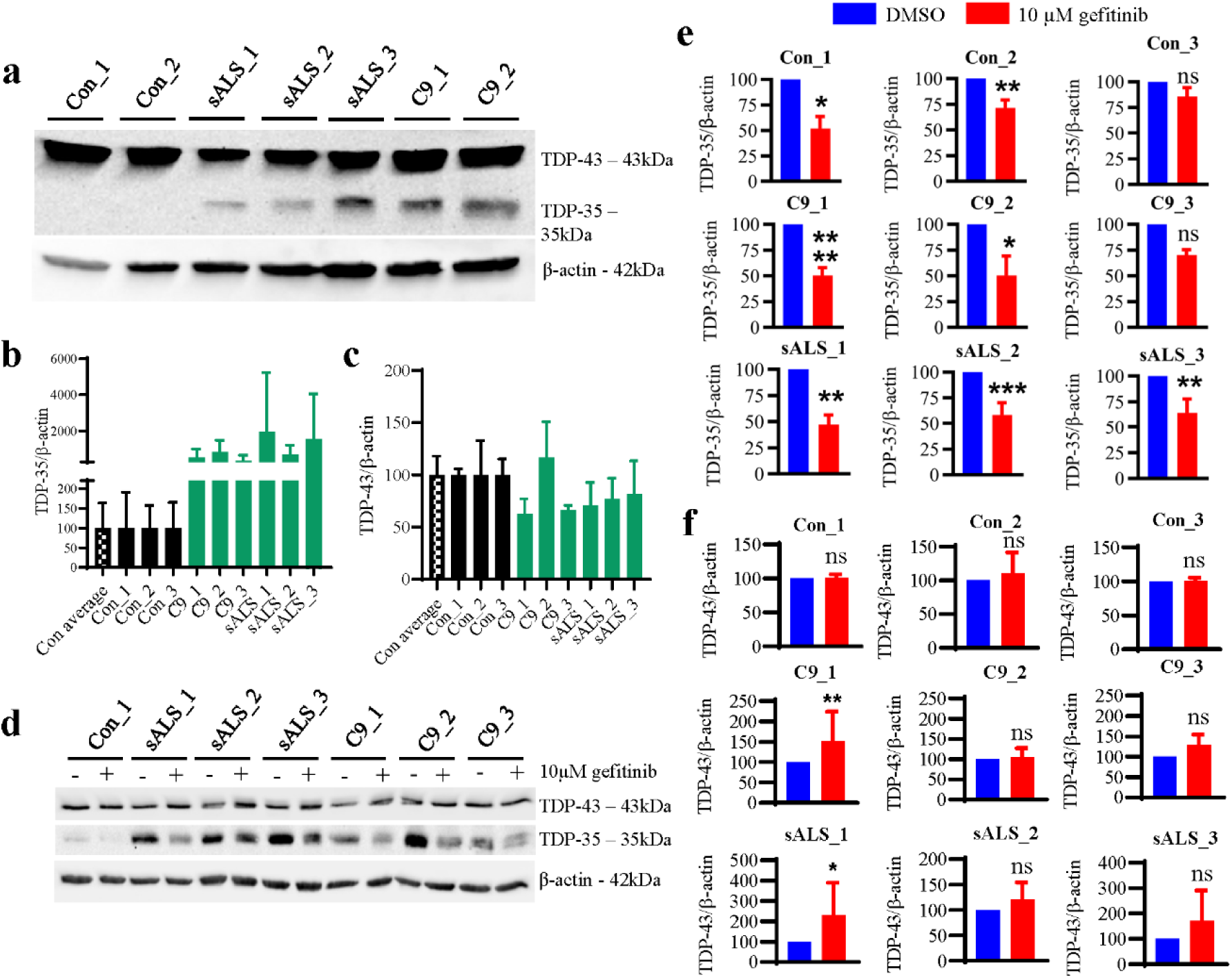
Gefitinib reduces 35 kDa fragment of TDP-43 in C9orf72-ALS and sALS iAstrocytes. **a)** Baseline expression of TDP-43 and TDP-35 cleavage product in C9orf72-ALS and sALS iAstrocytes and controls was assessed via Western blotting. **b)** Densitometry quantification of TDP-35 in iAstrocytes. Data normalised to control iAstrocytes = 100. Data = mean ± SD, n=3-6. One-way ANOVA with multiple comparisons test (vs. control average), ns. Data = mean ± SD, n=3-6. **c)** Densitometry quantification of TDP-43 in iAstrocytes. Data normalised to control iAstrocytes = 100. Data = mean ± SD, n=3-6. One-way ANOVA with multiple comparisons test (vs. control average), ns. **d)** Control, C9orf72 and sALS were treated with 10μM gefitinib and 0.1% (v/v) DMSO vehicle for 48h and the levels of TDP-43 and TDP-35 were assessed via Western blotting. **e)** Densitometry quantification of TDP-35 in iAstrocytes following a treatment with 10µM gefitinib. Data = mean ± SD, n = 3-10. Data normalised to DMSO = 100. Mann-Whitney test, *P < 0.05; **P < 0.01; ***P < 0.001; ****P<0.0001. **f)** Densitometry quantification of TDP-43 in iAstrocytes following a treatment with 10µM gefitinib. Data = mean ± SD, n = 3-7. Data normalised to DMSO = 100. Mann-Whitney test, *P < 0.05; **P < 0.01; ***P < 0.001; ****P<0.0001.

Following gefitinib treatment for 48 h to recapitulate the co-culture conditions, we observed a marked reduction in the levels of TDP-35 in all patient samples with the exception of C9_3, where the decrease was not statistically significant (P = 0.1; Fig. 3D-E). Interestingly, iAstrocytes from healthy individuals, which displayed very low levels of TDP-35 at baseline, potentially due to culture-induced stress, showed a further decrease after gefitinib treatment (Fig. 3E). Levels of the full-length TDP-43, on the other hand, were not affected consistently by gefitinib treatment, as we observed a significant increase only in C9_1 (P = 0.002) and sALS_1 (P = 0.029), whilst the remaining ALS lines, which display lower TDP-43 levels, showed no significant increase (Fig. 3F). We explored further whether gefitinib specifically attenuated TDP-43 proteinopathy by reducing cytoplasmic levels of TDP-35, whilst increasing full length TDP-43 in the nucleus. We conducted subcellular nucleocytoplasmic fractionations on iAstrocytes, confirmed by enrichment of the nuclear histone chaperone SSRP1 in the nuclear fraction and enrichment of the microtubule marker Tuj1 in the cytoplasmic fraction (Supplementary Fig. S2A). Following gefitinib treatment, levels of full length TDP-43 and TDP-35 were significantly reduced in the cytoplasmic fraction of C9_1 iAstrocytes (P = 0.012, P = 0.001 respectively; Fig. S2B). Gefitinib treatment also led to a reduction in TDP-35 in the nuclear fraction (P = 0.02; Fig. S2C) but did not significantly increase the levels of full-length TDP-43 in the nucleus. Consistently with the whole cell lysate, the fractionation data confirmed that gefitinib effect on full length TDP-43 levels cannot be attributed to re-localisation of TDP-43 to the nucleus. All together, these data confirm that gefitinib reduces TDP-43 proteinopathy and specifically highlights that the treatment reduces cytoplasmic and nuclear levels of TDP-35.

Since motor neurons are the primary cell type degenerating in ALS and they display the highest levels of proteinopathy in post-mortem tissues (14), we assessed the effect of gefitinib also on this cell type. iNPC-derived induced neurons (iNeurons) obtained from C9_1 and sALS_1 confirmed the data obtained in iAstrocytes by showing a decrease in TDP-35 levels following treatment with gefitinib (Fig. S3A, C), without an increase in TDP-43 (Fig. S3B), thus confirming gefitinib’s ability to reduce TDP-43 proteinopathy also in neurons.

These results prompted us to test whether longer gefitinib treatments would lead to a further reduction of TDP-43 proteinopathy, hence we treated patient iAstrocytes for 48 h, 72 h and 120 h with gefitinib before collecting the cells for immunoblotting (Fig. S4A). Levels of TDP-43 remained unaffected in all treatment conditions (Fig. S4B). The data confirmed the previously observed reduction in TDP-35 after 48 h alone but no further reduction in TDP-35 was achieved by longer gefitinib treatments (Fig. S4C).

Although EGFR knock-down in iAstrocytes did not improve MN survival in the co-culture, we tested whether EGFR is involved in mediating the TDP-35 reduction. As before, EGFR expression was knocked-down by using an adenonovirus expressing a validated shRNA and RFP or RFP only for 72 h, and cells were harvested for immunoblotting (Fig. S5A). We observed no change in TDP-43 (Fig. S5B) and TDP-35 (Fig. S5C) in iAstrocytes where EGFR levels were knocked-down. Interestingly, and consistently with this result, treatment with erlotinib, another potent first-generation EGFR inhibitor used to treat NSCLC, also did not lead to MN rescue in co-culture (Fig. 2C) or a decrease in TDP-43 proteinopathy (Fig. S6A-C), further suggesting that EGFR modulation is not key for gefitinib to rescue MN survival and target TDP-43 proteinopathy.

### Gefitinib induces autophagy in healthy cells and ALS iAstrocytes

Since gefitinib treatment reduced the levels of TDP-35 without a consistent effect on full-length TDP-43, we hypothesised that the decrease in TDP-35 is mediated via upregulation of autophagy rather than reduction in full-length protein cleavage or nuclear relocalisation. First, we validated gefitinib’s ability to activate autophagy in the human embryonic kidney (HEK293) cell line (Fig. 4A). Cells were treated for 6 h with bafilomycin A1, which inhibits lysosome-mediated cargo degradation by blocking the fusion of lysosomes with autophagophores (56), either alone or in combination with gefitinib. An mTORC1 inhibitor, rapamycin, was used as a positive control for autophagy activation. Levels of LC3-II increased by 1.38-fold in cells treated with gefitinib + bafilomycin A1 compared to bafilomycin A1 alone (P = 0.035; Fig. 4B), indicating autophagy activation. Consistently, the same treatment resulted in a 1.42-fold increase in LC3-II/LC3-I ratio when the two treatments were compared (P = 0.024; Fig 4C).

**Figure 4.**
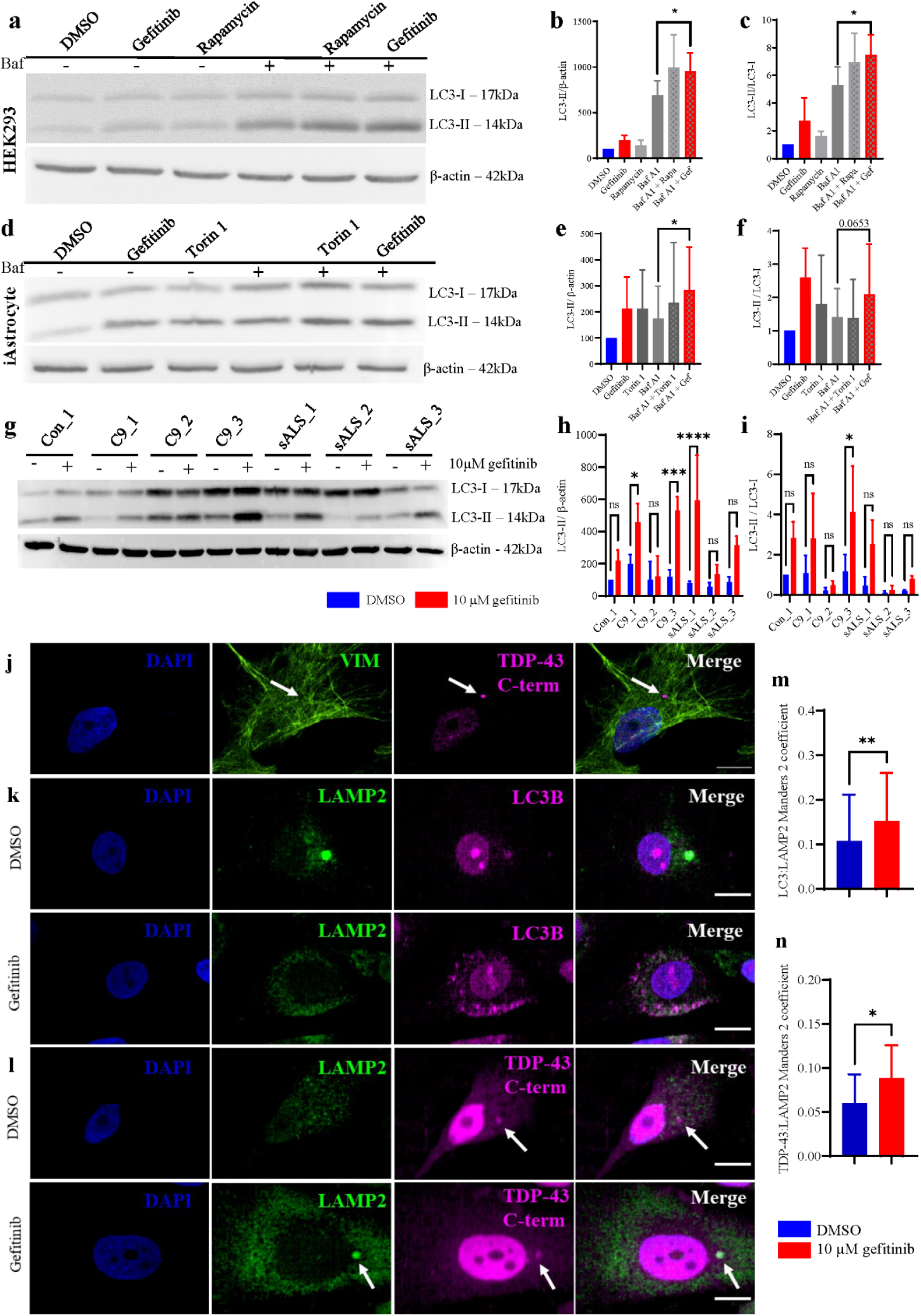
Gefitinib degrades TDP-43 35 kDa fragment via autophagy. **a)** Healthy HEK293 cells were treated for 6h with 0.1% (v/v) DMSO vehicle control, 10 µM gefitinib, 500 nM rapamycin, with and without 100 nM bafilomycin A1 (Baf) co-treatment. The induction of autophagic flux, as indicated by LC3-II levels and LC3-II/LC3-I ratio, was subsequently measured via western blotting. **b)** Densitometry quantification of LC3-II – LC3-I ratio. Data normalised to DMSO vehicle = 1. Data = mean ± SD; n = 5. Paired t-test (Baf A1 vs Baf A1 + Gef), *P < 0.05. **c)** Densitometry quantification of LC3-II expression. Data normalised to DMSO vehicle = 100. Data = mean ± SD; n = 5. Paired t-test (Baf A1 vs Baf A1 + Gef), *P < 0.05. **d)** Healthy control iAstrocytes were treated for 16h with 0.1% (v/v) DMSO vehicle control (0.1% v/v), 10 µM gefitinib, 250 nM Torin 1, with and without 100 nM bafilomycin A1 co-treatment. The induction of autophagic flux, as indicated by LC3-II levels and LC3-II/LC3-I ratio, was subsequently measured via western blotting. **e)** Densitometry quantification of LC3-II expression. Data normalised to DMSO vehicle = 100. Data = mean ± SD; n=6 (Con_3 n=2, Con_4 n=2, Con_5 n=2). Paired t-test (Baf A1 vs Baf A1 + Gef), *P < 0.05. **f)** Densitometry quantification of LC3-II – LC3-I ratio. Data normalised to DMSO vehicle = 1. Data = mean ± SD; n=6 (Con_3 n=2, Con_4 n=2, Con_5 n=2). Paired t-test (Baf A1 vs Baf A1 + Gef), ns. **g)** Expression of LC3 in control, C9, and sALS iAstrocytes after treatment with DMSO 0.1% v/v or gefitinib 10 µM for 48h. **h)** Densitometry quantification of LC3-II expression. Data normalised to DMSO vehicle-treated Con_1 iAstrocytes = 100. Data = mean ± SD; n = 3 Two-way ANOVA with Šidák’s multiple comparison’s test, *P < 0.05; ***P < 0.001; ****P<0.0001. **i)** Densitometry quantification of LC3-II – LC3-I ratio. Data normalised to DMSO vehicle-treated Con_1 iAstrocytes = 1. Data = mean ± SD; n = 3-4. Two-way ANOVA with Šidák’s multiple comparison’s test, *P < 0.05. **j)** Example image of untreated C9_1 iAstrocyte line showing TDP-43 C-terminal cytoplasmic aggregate enclosed within a vimentin cage (white arrow). Scale = 20 μm. **k)** Representative images of C9_1 iAstrocytes immunostained for LC3B and LAMP2, following a 48h treatment with 0.1% (v/v) DMSO vehicle control or 10 µM gefitinib. Scale: 20 μm. **l)** Representative images of C9_1 iAstrocytes immunostained for TDP-43 C-terminal and LAMP2, following a 48h treatment with 0.1% (v/v) DMSO vehicle control or 10 µM gefitinib. Scale: 20 μm. **m)** Manders coefficient of TDP-43 co-localisation with LAMP2 was determined using JACoP plugin for Fiji for C9_1 iAstrocytes. Data = mean ± SD; n=3. Paired t-test, **P<0.005. **n)** Manders coefficient of TDP-43 co-localisation with LAMP2 was determined using JACoP plugin for Fiji for C9_1 iAstrocytes. Data = mean ± SD; n=3. Paired t-test, *P < 0.05.

Next, we repeated the autophagy assay in iAstrocytes derived from healthy controls (Fig. 4D). In these experiments, rapamycin was replaced with torin 1 due to its ability to inhibit mTORC1 and mTORC2 rather than mTORC1 only. Here, gefitinib + bafilomycin A1 led to a significant increase in LC3-II levels over 16 h (P = 0.033; Fig. 4E). LC3-II/LC3-I ratio also increased after the treatment, albeit did not reach statistical significance (P = 0.065; Fig. 4F).

Interestingly, bafilomycin A1 treatment, even at low concentrations (10–500 nM) induced high levels of toxicity in ALS iAstrocytes (data not shown). This made the assay unsuitable for assessment of gefitinib’s ability to activate autophagy in patient cells, which are known to display defects in this pathway (9). For this reason, we have measured the levels of LC3 after a 48-h treatment with 10 μM gefitinib (Fig. 4G), matching the conditions at which we have observed maximal levels of TDP-35 reduction. ALS iAstrocytes generally expressed higher levels of LC3-II after gefitinib treatment compared to vehicle, albeit there was variability between different cell lines (Fig. 4H). The LC3-II/LC3-I ratio followed a similar pattern of response (Fig. 4I). Overall, the results indicate that gefitinib is capable of inducing autophagy in ALS patient iAstrocytes.

To further validate these findings, we tested the effect of gefitinib on autophagy in the absence of autophagy inhibitors. We also assessed its ability to promote autolysosomal fusion, which could, in turn, lead to degradation of TDP-35. For this purpose, C9_1 patient and Con_1 control iAstrocyte line were treated for 48 h with 10 µM gefitinib or 0.1% (v/v) DMSO vehicle control and processed for immunocytochemistry. First, we verified the presence of cytoplasmic aggregates containing the C-terminal of TDP-43, which is a major component of TDP-35, within vimentin cages in C9_1 iAstrocytes at baseline using confocal microscopy (Fig. 4J). Cells were then stained for the autophagy marker LC3B, lysosomal membrane component LAMP2, and the C-terminal domain of TDP-43 (Fig. 4K, L). Co-localization of LAMP2 signal with LC3B and TDP-43 respectively was assessed using confocal imaging and quantified. Compared to their corresponding vehicle controls, gefitinib-treated ALS patient cells showed a significant increase in LC3 co-localization with LAMP2 (C9_1: 1.43-fold change, P = 0.0034; Fig. 4M), suggesting autolysosomal fusion. Likewise, significantly more TDP-43 signal overlapped with LAMP2 following gefitinib administration (C9_1: 1.55-fold change, P = 0.0119), suggesting an increased lysosomal degradation of the C-terminal fragments of TDP-43 (Fig. 4N).

### Gefitinib delays disease onset in SOD1^G93A^ mice

Given the neuroprotection achieved by gefitinib treatment in co-cultures with ALS iAstrocytes across different genotypes, including SOD1, we proceeded to test gefitinib for efficacy *in vivo* using the well-characterised SOD1^G93A^ mouse model of ALS. Although brain penetrant, gefitinib is a substrate for the CNS efflux transporters P-glycoprotein (P-gp) and breast cancer resistance protein (BCRP) at the blood brain barrier (BBB) (57), which results in poor brain exposure. Previously, mice dosed with gefitinib and co-dosed with elacridar ‒ a dual P-gp and BCRP inhibitor – had improved gefitinib brain exposure compared to mice dosed with gefitinib alone (57). Therefore, we tested a similar dosing regimen in SOD1^G93A^ mice (Fig. S7A, B). Animals were orally (P.O.) dosed with 100 mg/kg elacridar or vehicle. Four hours later, animals were dosed with either 100 or 200 mg/kg gefitinib, and tissue was collected after 2 h. In the groups of animals dosed with 100 mg/kg gefitinib, the 100 mg/kg elacridar pre-dose increased free gefitinib in the brain by 7-fold, from 11.3 ± 1.2 nM to 77.4 ± 22.5 nM (P = 0.165) (Fig. 5A). In the animals dosed with 200 mg/kg gefitinib, the elacridar pre-dose increased free gefitinib in the brain by 14-fold from 21.5 ± 3.5 nM to 309.5 ± 79.6 nM (P < 0.0001; Fig. 5A, Fig. S7D). As expected, elacridar treatment did not affect gefitinib levels in the blood (Fig. S7C, E). Therefore, we took forward the gefitinib and elacridar co-dosing regimen to test gefitinib for efficacy in the SOD1^G93A^ mice (Fig. S8A). SOD1^G93A^ mice were dosed P.O. daily with either the vehicle control, 200 mg/kg gefitinib only, 100 mg/kg elacridar only, or 200 mg/kg gefitinib and 100 mg/kg elacridar from 25 days of age. As measured by the presence of the hind limb splay defect and hind limb tremor, neurological symptom onset was delayed from 66.6 ± 5.9 to 74.1 ± 3.4 days old (P = 0.002) in both the gefitinib + elacridar treated animals compared to elacridar-only treated animals (Fig. 5B, Fig. S8B). There was no significant difference in onset between the vehicle control group and gefitinib only dosed group of animals (Fig. 5B).

**Figure 5.**
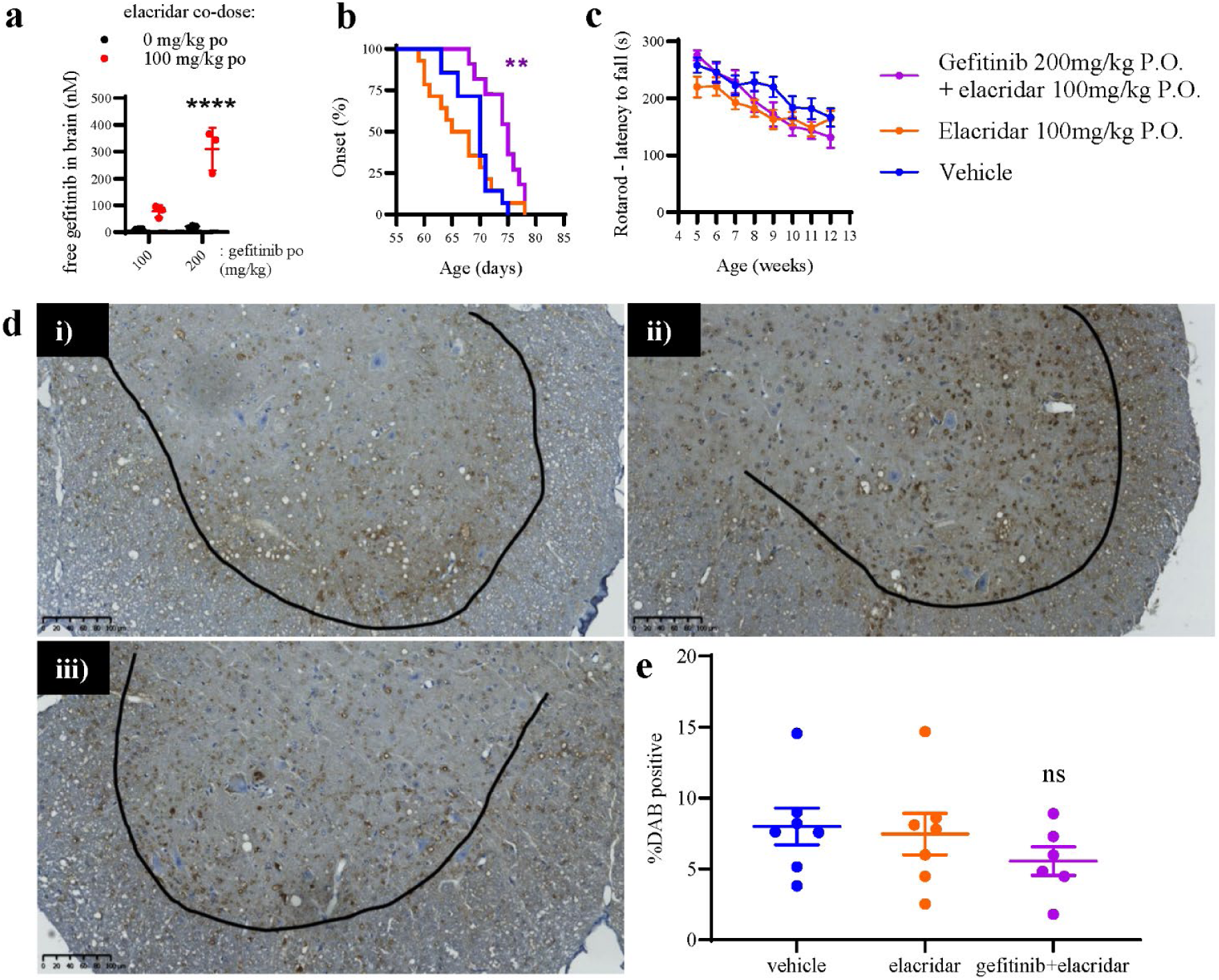
High brain exposure of gefitinib delays neurological symptom onset in SOD1^G93A^ mice, but does not rescue misfolded SOD1 pathology. **a)** Female SOD1^G93A^ C57BL/6 mice were either orally dosed with 100 mg/kg elacridar (-4 h), or untreated (no dose control group). Mice were then given a single oral dose of gefitinib at either 100 mg/kg po or 200 mg/kg po. Blood and brain were collected 2 h after gefitinib administration, and gefitinib levels were measured in brain using UHPLC - TOF mass spectrometry. Free gefitinib in the brain was calculated using total gefitinib brain levels and gefitinib brain tissue binding. n=3. Individual data points, as well as mean ± SD are shown. Two-way ANOVA with Šidák’s multiple comparisons; ****P < 0.0001. **b)** Female SOD1^G93A^ mice were dosed daily from 25 days of age with vehicle control P.O. (blue), 200 mg/kg gefitinib P.O. (green), 100 mg/kg elacridar P.O. (orange), or 200 mg/kg gefitinib and 100 mg/kg elacridar P.O. (purple). Neurological symptom onset was defined when both a hind limb splay defect and hind limb tremor were observed in an animal, and the percentage of animals with onset as a function of age is plotted on a Kaplan-Meier curve. n = 11-14. One-way ANOVA with Tukey’s post-hoc test; **P < 0.01. **c)** Rotarod performance assessed once a week from 5 weeks of age and was measured as time to fall in seconds. Two-way ANOVA with Tukey’s post-hoc test. **d)** Representative images of misfolded SOD1-stained spinal cords of SOD1^G93A^ mice treated with drug vehicle (i), elacridar (ii), or elacridar+gefitinib (iii). Scale bar: 100μm. **e)** Quantification of mSOD1-positive area (% DAB positive area). Data = mean ± SEM; n = 6-7. One-way ANOVA with Dunnett’s post-hoc test; all comparisons ns.

Rotarod (Fig. 5C, Fig. S8C) and running wheel (Fig. S8D, E) motor performance measures did not reveal a significant improvement in the gefitinib treated groups. On the contrary, time and distance run per day on the running wheels were both reduced in mice dosed with gefitinib alone or elacridar only, and this was further reduced in the gefitinib plus elacridar dosed animals (Fig. S8D, E). This correlated with a decrease in weight (Fig. S8F) and lethargic behaviour, which is a common side effect of anti-cancer drugs such as gefitinib.

To explain the observed shift in the symptom onset, markers of misfolded SOD1 (mSOD1) pathology and neuroinflammation were assessed. Here, we hypothesised that gefitinib-mediated autophagy activation might have an effect on the aggregates of mSOD1, similar to TDP-43 in iAstrocytes *in vitro*. Mice were sacrificed at 90 days, and spinal cords were stained for mSOD1 (Fig. 5D), astrocyte marker GFAP, and IBA-1 microglial marker (Fig. S9A). Spinal cord lysates were used to verify gefitinib’s engagement with its primary target EGFR, however, no signal was detected (data not shown). We observed a consistent, albeit not statistically significant, trend for a reduction of mSOD1 in animals treated with gefitinib + elacridar (Fig. 5E, Fig. S9D). There was no significant difference in GFAP or IBA-1 staining intensity between any of the treatment groups (Fig. S9B, C).

## Discussion

ALS is a devastating, complex disorder of the motor system, characterised by fast progression and poor prognosis. With no disease-course modifying drugs available, there is a critical need for new effective therapies for ALS, as the two currently available drugs, riluzole and edaravone, offer limited improvements to patients’ health (58,59). The process of drug discovery for ALS has been marred by failures (1, 60), further highlighting the need for novel approaches. Lack of knowledge of the mode of action of a compound and lack of appreciation of the heterogeneity of ALS are amongst the main overlooked factors leading to failures of phase II clinical trials (1, 61, 62).

The application of AI to drug repurposing has the potential to significantly accelerate the discovery of therapeutics at a much-reduced cost so we used high-dimensional structured and unstructured data mining in a search for drugs with repurposing potential for ALS. We tested the five drugs identified in our AI workflows in our iNPC model which has the advantage of retaining ageing features (49), and which allowed us to study TDP-43 proteinopathy in iAstrocytes at baseline without a need to further stress the cells. This contrast with commonly utilised models where TDP-43 has been overexpressed or artificially truncated (19, 63–68). Although MNs are the main cell type that perishes in ALS, the disease has been long described as a non-cell autonomous disease (69, 70). Improving the health of astrocytes, which exhibit a loss of their supportive role in ALS patients, can consequently improve the prognosis and/or prolong the survival of the MNs themselves.

Of the five identified drugs, gefitinib showed the most consistent protection of MNs, associated with a significant reduction in fragmented TDP-35 in the cytoplasm of most iAstrocytes. While mutant full length cytoplasmic TDP-43 is more toxic than its fragments (due to its longer half-life (18)), TDP-35 and other C-terminal-containing fragments are the main drivers of aggregation through directly binding and sequestering the RNA-recognition motif 1 (RRM1) of TDP-43 (71,72). Based on these findings, several papers have shown that elimination of TDP-43 aggregates is beneficial for cell survival (26, 73–75). Accordingly, the gefitinib-mediated reduction in cytoplasmic TDP-35 may be the mechanism underlying the protection of MNs from the effects of the toxic ALS iAstrocytes.

We then examined the most likely mechanism suggested by the AI workflow by which gefitinib reduced the cytoplasmic levels of TDP-43 and TDP-35, i.e. autophagy, which has been shown to be increased by gefitinib in several cancer cell lines (76, 77). Gefitinib has been previously extensively described as an autophagy activator (76, 78), especially in the context of EGFR activation, which causes autophagy suppression by binding Beclin 1 and blocking its activity (79).

Gefitinib-mediated inhibition of constitutively active EGFR blocks its kinase activity and has been reported to cause increase in autophagic flux through suppression of AKT and mTOR (76). Gefitinib’s ability to boost autophagy, however, has also been shown in cells expressing low levels of EGFR, such as mouse fibroblasts or cells where EGFR had been knocked out through CRISPR/Cas9 approaches (80). This indicates that gefitinib can activate autophagy through an EGFR-independent mechanism. Importantly, it is known that C9orf72 hexanucleotide expansion in ALS causes defects in autophagy initiation and results in lysosomal abnormalities, preventing normal autolysosomal fusion and thus conclusion of autophagy (81–83). Indeed, all ALS iAstrocytes lines in this study showed a baseline accumulation of LC3-I, which is emblematic of an autophagy dysfunction (84). It is interesting that bafilomycin A1 caused a high degree of toxicity selectively in ALS cells as opposed to controls. This might be due to an increase in sensitivity of these cells to the pathway inhibitor, due to the already existing impairment in the pathway. To overcome these limitations, we have demonstrated an increased co-localization of LC3 and LAMP2 after gefitinib treatment, suggestive of autolysosomal fusion in process. Moreover, we have observed an increased localization of TDP-43 C-terminal into lysosomes, further indicating that gefitinib promotes autophagic degradation of cytoplasmic TDP-43 and its cleavage products.

Although appealing, few repurposing drug candidates have reasonable CNS penetrance. The neuroprotective action of gefitinib in co-cultures with SOD1 patient iAstrocytes, lead us to test this drug further in SOD1^G93A^ mice, as we hypothesised that activation of autophagy would limit neurodegeneration in this model. We designed an approach to maximise CNS exposure by inhibiting the PGP/BCRP transporters known to drive efflux of gefitinib from the CNS, which gave maximum probability of target engagement. Other drugs with repurposing potential might need to be structurally redesigned to allow for brain penetrance, thus potentially dampening the advantage of a repurposing strategy.

Co-dosing of gefitinib and elacridar was, indeed, more effective than gefitinib treatment alone and led to a significant shift in disease onset accompanied by a non-significant trend in reduction of misfolded SOD1 in the spinal cord. These results are in line with the finding that protecting MNs results in a delay in disease onset (85). It is possible that activation of autophagy in both astrocytes and neurons is responsible for the beneficial effects observed in this study. Progressive motor symptoms, however, did not improve, likely due to unknown negative effects of gefitinib exposure in the CNS. The observed side effects, such as lethargy and weight loss, are also seen in NSCLC patients taking gefitinib and are consistent with on-target EGFR inhibition in peripheral organs (86) and could have led to reduced motor activity. Interestingly, a study investigating the EGFR inhibitor erlotinib in SOD1^G93A^ mice observed a similar delay in symptom onset, but without a rescue of MN survival at the NMJ (87). Erlotinib, previously described as a more potent inhibitor of EGFR than gefitinib (88), underperformed in the co-culture assay used here and had no effect on TDP-43 proteinopathy (Supplementary Fig. S3) in comparison with gefitinib. This indicates that gefitinib carries an additional neuroprotective property absent in erlotinib, which is independent of EGFR activity in our cell model. Based on our *in vitro* data, the additional protective mechanism is likely the activation of autophagy. The involvement of different kinases targeted by gefitinib, however, cannot be excluded. Indeed, lines of evidence suggest that targeting autophagy alone might not be sufficient to achieve neuroprotection (89), hence gefitinib’s neuroprotective effect *in vitro* and *in vivo* could be due to multiple targets and MOAs. Further studies could employ machine learning-driven knowledge graphing in order to shortlist those mechanisms.

Indeed, the ability to combine vast amounts of information and reported MOAs that are relevant across disease areas is one of the main advantages of using AI-powered drug repurposing approaches. In addition, the speed at which new hypotheses are generated is unprecedented and might offer a new and more effective avenue for drug discovery in ALS and other neurodegenerative disorders.

## Materials and Methods

### Conversion of skin fibroblasts to induced neural progenitor cells (iNPCs)

Skin fibroblasts from healthy donor controls and ALS patients were reprogrammed as previously described (47). Briefly, 10,000 fibroblasts were seeded into a well of a 6-well plate and incubated for 24 h in fibroblast media (DMEM (Lonza) and 10% FBS (Biosera)). Cells were treated with retroviral vectors containing OCT3/4, Sox2, KLF4 and C-MYC for 12 h. Cells were incubated for a further 24 h in fibroblast media. The cells were then washed in PBS and then the media was changed to NPC conversion medium (DMEM/F12 (1:1) GlutaMax (Gibco), 1% N-2 (Gibco, catalogue number: 11520536), 1% B-27 (Gibco, catalogue number: 11530536), 20 ng/mL FGF2 (Peprotech), 20 ng/mL EGF (Peprotech), and 5 ng/mL heparin (Sigma-Aldrich)). Cells were expanded into individual wells on a 6-well plate once cell morphology changed and cells developed a sphere-like form. The media was switched to NPC proliferation media (DMEM/F12 (1:1) GlutaMax, 1% N-2, 1% B-27, and 40 ng/mL FGF2) once an iNPC culture was established. Clinical information of the patients whose samples were used in this study are detailed in Table 1.

**Table 1.** Clinical information of donors from which iNPC lines were derived.

| Sample | Mutation | Sex | Ethnicity | Age at biopsy collection | Onset to death (months) |
| --- | --- | --- | --- | --- | --- |
| Con_1 | - | M | Caucasian | 40 | - |
| Con_2 | - | M | Caucasian | 55 | - |
| Con_3 | - | F | Caucasian | 64 | - |
| Con_4 | - | F | Caucasian | 52 | - |
| Con_5 | - | M | Caucasian | 31 | - |
| C9_1 | C9orf72 | M | Caucasian | 66 | 31.7 |
| C9_1 | C9orf72 | M | Caucasian | 50 | 27 |
| C9_3 | C9orf72 | F | Caucasian | 66 | 19.4 |
| sALS_1 | sALS | F | Caucasian | 61 | 21 |
| sALS_2 | sALS | M | Caucasian | 29 | 90 |
| sALS_3 | sALS | M | Caucasian | 47 | 72 |
| SOD1_1 | SOD1 (A4V) | F | Caucasian | 40 | - |

### Human iAstrocyte differentiation and maintenance

iAstrocytes were produced as previously described (47,48). Briefly, iNPCs were switched to iAstrocyte differentiation media (DMEM (Lonza, catalogue number: 12-741F), 10% FBS (Biosera, catalogue number: FB-1090), 0.2% N-2 (Gibco)), and differentiated for 7 d. Cells were grown in Fibronectin-coated 10 cm dishes.

### Murine Hb9-GFP+ MN differentiation

Murine Hb9-GFP+ MN were produced as previously described (48). Briefly, murine embryonic stem cells (mESC) containing GFP controlled by the MN-specific promoter Hb9 (kind gift from Thomas Jessel, Columbia University, New York) were differentiated to produce Hb9-GFP+ MN. Hb9-GFP mESC were maintained by culturing on mouse embryonic fibroblasts (Merk) in mESC media (KnockOut DMEM (Gibco), 15% (v/v) embryonic stem-cell FBS (Gibco, catalogue number: 11500526), 2 mM L-glutamine (Lonza, catalogue number: BE-17-605E), 1% (v/v) non-essential amino acids (Gibco, catalogue number: 12084947) and 0.00072% (v/v) 2-mercaptoethanol (Sigma, catalogue number: M3148)). Hb9-GP mESC were differentiated into MN via embryoid bodies (EBs). mESC were lifted using 1x trypsin (Lonza, catalogue number: BE02-007E) in PBS, resuspended in EB media (DMEM/F12 (Gibco), 10% (v/v) knockout serum replacement (Gibco), 1% N2 (Gibco), 1 mM L-glutamine, 0.5% (w/v) glucose (Sigma, catalogue number: G7021) and 0.0016% (v/v) 2-mercaptoethanol), and cultured in non-adherent petri dishes. The media was replenished daily. On days 2-7 of differentiation, 2 μM retinoic acid (Sigma, catalogue number: R2625) and 0.5 μM smoothened agonist (Sigma, catalogue number: 566660) were added to the EB media. At day 7, EBs were dissociated using papain (Sigma, catalogue number: P4762).

### iAstrocyte-Hb9-GFP+ MN co-culture assay

The 384-well plate co-culture assay using human iAstrocytes and murine Hb9-GFP+ MN was performed as described (48). At day 6, iAstrocytes were treated with ambroxol, erlotinib, gefitinib, RTA-408, and siramesine (all compounds from Cayman chemicals; catalogue numbers 19377, 10483, 13166, 17854, and 21817 respectively); the final DMSO (Sigma-Aldrich) concentration was 0.1% (v/v). 384-well co-culture plates were imaged on an In Cell Analyzer 2000. The number of viable GFP+ MN (GFP+ cell bodies with at least 1 neurite attached) were counted at 24 h and 72 h of co-culture using the Columbus Image Data Storage and Analysis System (PerkinElmer). % MN survival was calculated as the number of MN at 72 h as a percentage of number of MN at 24 h, and then normalised to the DMSO control value.

### HEK293 cell culture and drug treatment

HEK293 cells were cultured in DMEM supplemented with 10% (v/v) FBS, 50 units/mL penicillin/streptomycin and sodium pyruvate at 37°C/5% CO2. For the autophagy activity assay, HEK293 cells were plated into 6-well plates and cultured until they reached 80% confluence. Cells were treated with 10 µM gefitinib, 500 nM rapamycin (Cayman Chemicals), 100 nM bafilomycin A1 (Alfa Aesar), or a combination of bafilomycin A1 with rapamycin or gefitinib for 6 h. The final DMSO concentration was 0.1% (v/v).

### Immunocytochemistry

iAstrocytes were differentiated as described before. 96 h after seeding in the differentiation medium, iAstrocytes were accutased and replated into 2.5 µg/mL fibronectin-coated 96-well optical plates at a density of 8000 cells/well. 24 h after replating, the culture media was replaced with media containing 10 µM gefitinib and 0.1% (v/v) DMSO vehicle. After 48 h, all media was removed, cells washed with PBS, and fixed with 4% paraformaldehyde for 10 min at RT. Nonspecific binding was blocked for 1 h at RT with 5% (v/v) horse serum and 0.5% (v/v) Triton®-100 in PBS. Cells were incubated overnight at 4°C with chicken polyclonal anti-vimentin (1:1,000, Merck Millipore, catalogue number: AB5733), rabbit polyclonal anti-TDP-43 C-terminal (1:300, Proteintech, catalogue number: 12892-1-AP), rabbit polyclonal anti-LC3B (1:1,000, Novus Biologics, catalogue number: 2220), or mouse monoclonal anti-LAMP2 (1:200, Santa Cruz, clone H4B4, catalogue number: sc-18822). Following an incubation with primary antibodies, cells were washed for 5 minutes with PBS containing 0.2% (v/v) Tween®-20 whilst placed on a shaker at 50 rpm, followed by two more washes in PBS. Cells were incubated with PBS-diluted secondary antibodies Alexa Fluor goat anti–chicken 488 IgG (H+L) (1:1,000, Invitrogen, catalogue number: A-11039), Alexa Fluor donkey anti–mouse 568 IgG (H+L) (1:1,000, Invitrogen, catalogue number: A-10037), Alexa Fluor goat anti–mouse 647 IgG (H+L) (1:1,000, Invitrogen, catalogue number: A-21235) for 1h at RT. Hoescht stain solution diluted 1:10,000 in PBS was added for 5 min at RT to label nuclei, followed by two 5-min washes in PBS. Stained plates were imaged using Opera Phenix™ High Content Screening System (Perkin Elmer) and analysed using associated Columbus Data Storage and Analysis System (Perkin Elmer).

### Immunohistochemistry

Collected tissue that has been embedded in wax on cover slips was rehydrated by incubation at RT in xylene twice for 5 min, 100% ethanol for 5 min, 95% ethanol for 5 minutes, 70% ethanol for 5 minutes and water for 5 minutes. For antigen retrieval, slides were incubated for 30 min at pressure level 0 in a pressure cooker. Following the antigen retrieval, slides were washed gently with water to cool the slides, after which tissue was washed in PBS on a shaker for 5 min at RT. Slides were then dried and tissue blocked for 20 min at RT in 5% BSA in PBS with 0.25% Triton X-100 to permeabilised the tissue. Following the block, the slides were dried, placed in a slide tray containing water to prevent antibody evaporation, and tissue incubated overnight at 4°C with chicken polyclonal anti-GFAP (1:500, Abcam, catalogue number: ab4674), or rabbit polyclonal anti-Iba1 (1:500, GeneTex, catalogue number: GTX1000042), all diluted in 1% BSA in PBS with 0.25% Triton X-100. The following day, slides were washed quickly six times with PBS and then three times for 8 min in PBS on a shaker. For fluorescent staining, slides were dried, blocked with 5% BSA for 10 min, dried again, and incubated in 1% BSA-diluted secondary antibodies Alexa Fluor goat anti–chicken 488 IgG (H+L) (1:1,000, Invitrogen, catalogue number: A-11039) and Alexa Fluor goat anti-rabbit 555 IgG (Heavy Chain) (1:1,000, Thermo Fisher, catalogue number: A27039). Slides were shielded from light and washed quickly twice with PBS, and then four times for 8 min each wash on a shaker. After a 5-min wash in distilled water, the slides were dried and a few drops of VECTASHIELD® HardSet™ mounting medium with DAPI (Vector Laboratories, catalogue number: H-1500-10) were applied to the tissue. A cover slip was lowered slowly onto each slide, and the mounting medium covered the slide evenly. Slides were left to dry at RT in the dark and stored at 4°C.

For DAB staining, following the blocking, slides were incubated overnight at 4°C with mouse anti-misfolded SOD1 (B8H10) (1:1000, MediMabs, catalogue number: MM-0070-P) and washed the following day as described above. The ABC-HRP complex (Vector Labs, catalogue number: PK6102) was prepared at least 30 minutes prior to use. Sections were washed twice in TBS for 5 minutes each wash and incubated in the DAB substrate (Vector Labs, catalogue number: SK4100). The sections were counterstained with hematoxylin gill 3 (TCS Biosciences, catalogue number: HS345-1L) for 2 minutes, rinsed with tap water and differentiated in 0.3% acid alcohol (3% hydrochloric acid in 95% ethanol). Sections were washed with Scott’s Tap Water (Leica, catalogue number: 3802900) to blue the hematoxylin and dehydrated through graded series of alcohols (70% ethanol, 95% ethanol, and two changes of absolute ethanol, for 1 minute each). Sections were cleared in xylene for 5 minutes and mounted in DPX mounting medium (Cell Path, catalogue number: SEA-1304-00A), then dried overnight at 40°C.

### Immunoblotting

iAstrocytes were seeded onto 6-well plates after 4 days of differentiation. Briefly, 1 mL of 2.5 µg/mL fibronectin (Sigma) was added to each well of the 6-well plates (Greiner-Bio, catalogue number: 781091); plates were incubated at RT for 5 min to coat. The 2.5 µg/mL fibronectin was removed, and 100,000-200,000 iAstrocytes in 2 mL iAstrocyte differentiation media were added to each well of the fibronectin-coated 6-well plates. If iAstrocytes were drug treated, media was removed 24 h after seeding, and 2 mL iAstrocyte differentiation media containing drug and 0.1% (v/v) DMSO was added per well. iAstrocytes were grown on 6-well plates until 7 days of differentiation and 100% confluence. Media was removed, and iAstrocytes were washed in PBS. iAstrocytes were lysed in lysis buffer (150mM NaCl, Protease Inhibitor Cocktail EDTA–free (Roche; used according to the manufacturer’s instructions)) on ice for 20 min. Lysates were centrifuged at 17,000 x g at 4°C for 5 min to clarify. Supernatant was incubated with laemmli buffer at 95°C for 5 min. 25 µg protein from iAstrocyte lysates were separated on 12% Tris-glycine SDS-polyacrylamide gels (SDS-PAGE) and transferred to Amersham Protran Premium 0.2 nitrocellulose membrane (GE Healthcare). Membranes were blocked in 5% milk (Marvel) in TBS-T at RT for 1 h. Membranes were then probed with rabbit polyclonal anti-TDP-43 C-terminal (1:1,000, Proteintech, catalogue number: 12892-1-AP), rabbit polyclonal anti-LC3B (1:1,000, Novus Biologics, catalogue number: 2220), mouse monoclonal anti-p62 lck ligand (1:800, BD Biosciences, clone 3/P62 LCK LIGAND, catalogue number: 610833), or mouse monoclonal anti-β-actin (1:10,000, Abcam, clone AC-15, catalogue number: ab6276), in 5% Milk/TBS-T at RT for 1h. Membranes were washed three times in TBS-T, then probed with goat polyclonal anti-rabbit-IgG HRP (1:5,000, DAKO, catalogue number: P0448) or goat polyclonal anti-mouse-IgG HRP (1:5,000, DAKO, catalogue number: P0447) in 5% Milk/TBT at RT for 1 h. Membranes were washed three times in TBS-T. The secondary antibodies were detected on the membranes using EZ ECL chemiluminescence detection kit for HRP (Geneflow Ltd) according to the manufacturer’s instructions, and the chemiluminescence was imaged using a G-BOX (Syngene). Densitometry analysis was performed using GeneTools software (Syngene, v4.3.10).

### Nucleocytoplasmic fractionation

iAstrocytes treated for 48 h with 10 µM gefitinib and 0.1% (v/v) DMSO were washed with PBS and incubated for 2-3 min in accutase at 37°C. Detached cells were collected in PBS using a trimmed P1000 pipette tip to avoid shearing the cells. Cells were centrifuged at 400 x g at 4°C for 4 min. After the supernatant was discarded, cells were washed briefly in a hypotonic lysis buffer (10 mM HEPES [pH 7.9], 1.5 mM MgCl_2_, 10 mM KCl, 0.5 mM DTT, Protease Inhibitor Cocktail EDTA–free) and the cell pellet lysed in a fresh hypotonic lysis buffer using a trimmed P1000 pipette tip. After a homogenous cell suspension was achieved, cells were passed through a 19g needle multiple times. Samples were incubated on ice for 15 min and centrifuged at 1,500 x g at 4°C for 3 minutes to pellet the nuclear cell compartment. Supernatant was transferred to a new tube and centrifuged at 3,500 x g at 4°C for 8 min, whilst the pellet was kept on ice in hypotonic lysis buffer. The supernatant was collected and transferred to a fresh tube to be centrifuged again at 17,000 x g at 4°C for 1 min. The resulting supernatant constituted the cytoplasmic cell fraction. The nuclear fraction pellet collected at the earlier step was resuspended in hypotonic lysis buffer using a trimmed P1000 pipette tip and centrifuged at 1,500 x g at 4°C for 3 min. This step was repeated up to 3 times to remove any remaining cytoplasmic fraction. The final pellet was resuspended in reporter lysis buffer (Promega, catalogue number: E4030, supplemented with Protease Inhibitor Cocktail EDTA–free) and passed multiple times through a 25g needle. After a 10-min incubation on ice, the cell suspension was centrifuged at 17,000 x g at 4°C for 5 min. The supernatant was transferred to a fresh tube and constituted the nuclear cell fraction. Both the nuclear and cytoplasmic cells fractions were mixed with 4X laemmli buffer and the resulting samples were then separated on SDS-PAGE gels as described above and the resulting nitrocellulose membranes were probed with rabbit polyclonal anti-TDP-43 C-terminal (1:1,000, Proteintech, catalogue number: 12892-1-AP), chicken polyclonal anti-Tuj1 (1:1,000, Merck Millipore, catalogue number: AB9354) and mouse monoclonal anti-SSRP1 (1:500, Abcam, clone 10D7, catalogue number: ab26212), followed by a HRP-conjugated secondary antibody incubation and imaging approaches described above.

### Transgenic C57BL/6 SOD1^G93A^ mouse model of ALS

Mice were originally obtained from the Jackson Laboratory, B6SJL-Tg (SOD1^G93A^)1Gur/J (stock number 002726) and were subsequently backcrossed onto the C57BL/6 background (Harlan UK, C57BL/6 J OlaHsd) for >20 generations to create a line on an inbred genetic background. The SOD1^G93A^ transgene was maintained as a hemizygous trait by breeding hemizygous males with wild-type females (C57BL/6J OlaHsd, Harlan UK). Hemizygous males for breeding were available from our facility upon request. Genotyping was performed as previously described (90).

### Oral dosing of SOD1^G93A^ mice

For the pharmacokinetic study 8-week-old female SOD1^G93A^ mice were dosed at 10 mL/kg. Elacridar (Sigma-Aldrich, catalogue number: SML0486) and gefitinib (Carbosynth, catalogue number: FG10816) were suspended in the vehicle: 0.5% (w/v) methylcellulose (Sigma-Aldrich, catalogue number: M0262) in water. Mice were weighed before the first dose. Mice were dosed with either 100 mg/kg elacridar P.O., or not dosed at all. Mice were dosed with either 100 mg/kg or 200 mg/kg gefitinib P.O.; the gefitinib dose was 4 h after the elacridar dose (if applicable). Mice were euthanised by intraperitoneal injection of 25 mL/kg pentobarbital 2 h after the gefitinib dose. For the gefitinib efficacy study, female SOD1^G93A^ mice were dosed once per day at 10 mL/kg from 25 days of age. Mice were dosed with either 0.5% (w/v) methylcellulose in water (vehicle control) P.O., 200 mg/kg gefitinib P.O., 100 mg/kg elacridar P.O., or 200 mg/kg gefitinib and 100 mg/kg elacridar P.O. Mice were weighed before each dose. Mice were singly housed in cages containing running fast-trac wheels. At day 90, mice were euthanised by intraperitoneal injection of 25 mL/kg pentobarbital.

### Behavioural testing of SOD1^G93A^ mice

Tremor and hind-limb splay were measured 3 times per week from 60 days of age. Neurological symptom onset was defined as the point when both enhanced tremor (score >0) and deficits in hind-limb splay (score >0) were scored. Tremor score was measured by suspending the mouse by the tail; forelimb and hind-limb tremor were recorded separately. The scores were: 0 = normal, 1 = mild tremor, 2 = moderate tremor, 3 = severe tremor. Hindlimb splay defects were recorded at the same time; left and right hind-limbs were recorded separately. The scores were: 0 = normal, 1 = mild splay defect, 2 = moderate splay defect, 3 = strong splay defect.

Rotarod testing was conducted once per week from 35 days of age. Rotarod training was performed on 3 consecutive days with 2 trials per day. Subsequently, mice were tested twice per day, with a rest period in between tests. The best score of the 2 tests was used for analysis. In each test (including training) the rotarod was accelerated from 4 to 40 rpm in 300 s. Latency to fall (s) was recorded for each mouse.

To measure running wheel performance, a magnet was fixed to a Fast-trac wheel (Bio-Serv), which was mounted on a Mouse Igloo (Bio-Serv). A Velo wireless 7 computer (Cateye) was also fixed to the outside of the running wheel cage, such that the computer measured the number and frequency of the Fast-Trac rotations. Time and distance run on the running wheel was measured every day for each mouse.

### Statistical analysis

Statistical details of experiments, including sample sizes and statistical tests used, are described in individual figure legends. P values of 0.05 and above were considered statistically significant. Unless otherwise stated, results are presented as mean ± SEM. Outlier analyses were performed prior to statistical analyses and outlier removed where necessary. Mean values were compared using a one-way ANOVA with Tukey’s multiple comparisons test. Mean values where two independent variables were present were compared using a two-way ANOVA with Šidák’s multiple comparison’s test. Comparisons of two groups were performed using a Student’s t-test, paired or unpaired, context-depending. Comparisons of two groups, where one value was set to 1 or 100%, were performed using a Mann-Whitney test or Wilcoxon matched-pairs signed rank test, context-depending. In experiments where two or more mean values were compared to a control value set to 1 or 100%, a Kruskal-Wallis test with Dunn’s multiple comparison’s test was used.

## Acknowledgments

We extend our deep gratitude to patients and healthy individuals who have donated the biomaterial used in this study. We thank Prof. Thomas Jessel for providing the Hb9-GFP+ stem cells. We also thank Ian Colidcott for his technical assistance with *in vivo* work conducted for this study, Daniel Fillingham and Lynne Baxter for technical assistance with mouse tissue histology, and Dr Adrian Higginbottom for drug screening facility assistance.

