## Supplementary figures for "Artificial intelligence-augmented drug discovery identifies gefitinib as a potential treatment for ALS"

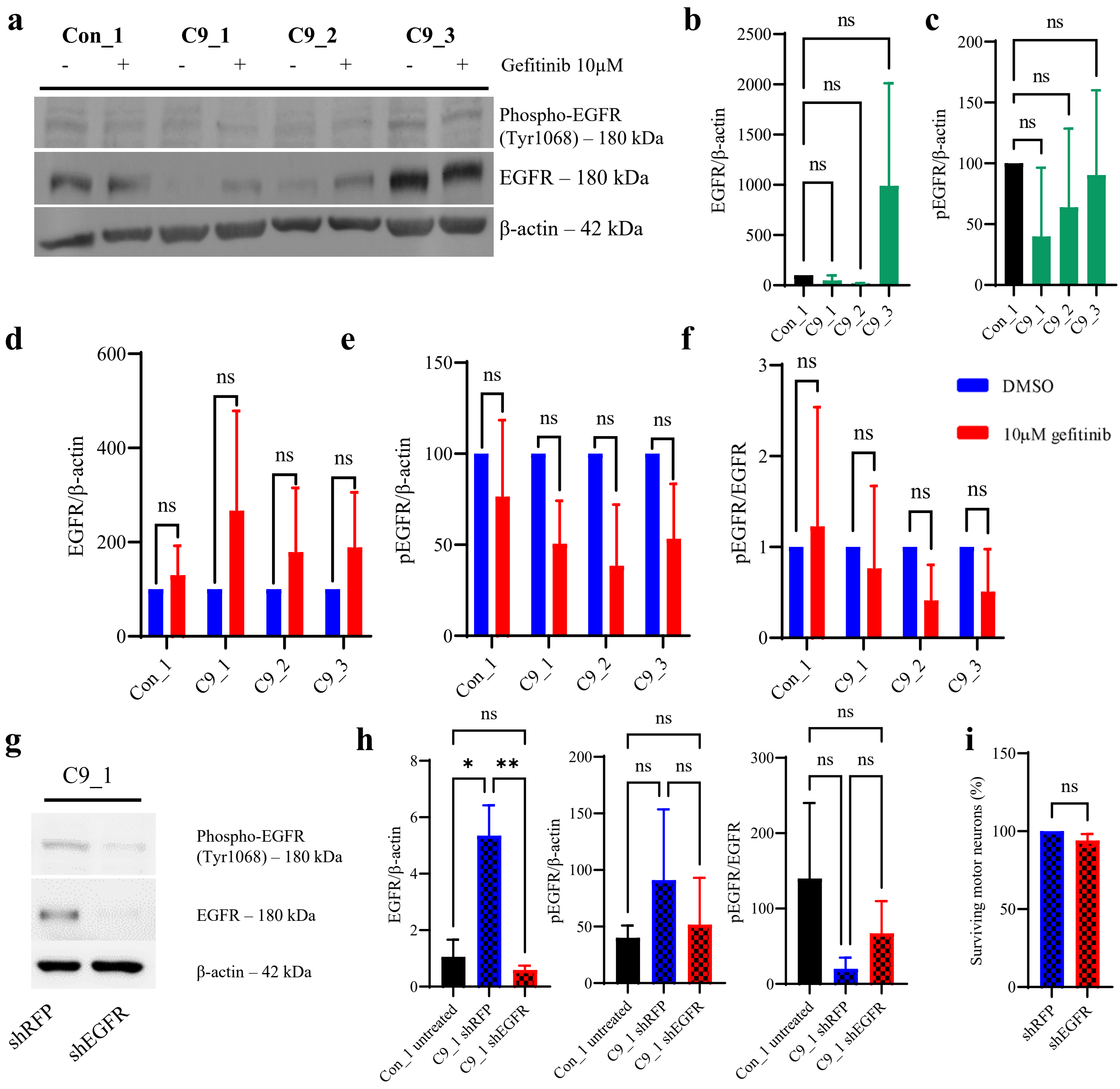


**Figure S1. Expression of EGFR in C9orf72 iAstrocytes. A)** Representative Western blot showing the expression level of EGFR and pEGFR (Tyr1068) in control and C9orf72 iAstrocytes before and after treatment with gefitinib. **B)** Densitometry quantification of the baseline expression levels of EGFR in Con_1 and C9orf72 iAstrocyte lines. Data = mean ± SD, n=3-4. Data normalised to Con_1 = 100. Kruskal-Wallis test (**P<0.005) with Dunn's multiple comparisons test, all comparisons ns. **C)** Densitometry quantification of the baseline expression levels of pEGFR (Tyr1068) in Con_1 and C9orf72 iAstrocyte lines. Data = mean ± SD, n=3-4. Data normalised to Con_1 = 100. Kruskal-Wallis test with Dunn's multiple comparisons test, all comparisons ns. **D)** Densitometry quantification of EGFR in iAstrocytes following a treatment with 10µM gefitinib. Data = mean ± SD, n=3-4. Data normalised to DMSO-treated iAstrocytes = 100. Mann-Whitney test, all comparisons ns. **E)** Densitometry quantification of pEGFR (Tyr1068) in iAstrocytes following a treatment with 10µM gefitinib. Data = mean ± SD, n=3-4. Data normalised to DMSO-treated iAstrocytes = 100. Mann-Whitney test, all comparisons ns. **F)** Analysis of pEGFR/EGFR ratio. Data = mean ± SD, n=3-4. Data normalised to DMSO-treated iAstrocytes = 100. Mann-Whitney test, all comparisons ns. **G)** Representative Western blot showing the expression level of EGFR and pEGFR (Tyr1068) in iAstrocyte line C9_1 following a knockdown with shRNA. **H)** Densitometry quantification of EGFR, pEGFR and pEGFR/EGFR ratio after EGFR knockdown in C9_1 iAstrocyte line. Data = mean ± SD, n=3. One-way ANOVA, *P<0.05, **P<0.005. **I)** iAstrocyte lines treated with shEGFR were co-cultured with murine Hb9-GFP+ MN. % MN survival was calculated as number of MN at 72h as a percentage of number of MN at 24h, and then normalised to the average of DMSO control value. Data = mean ± SD, n=4. Data normalised to DMSO = 100. Wilcoxon matched-pairs signed rank test, ns.

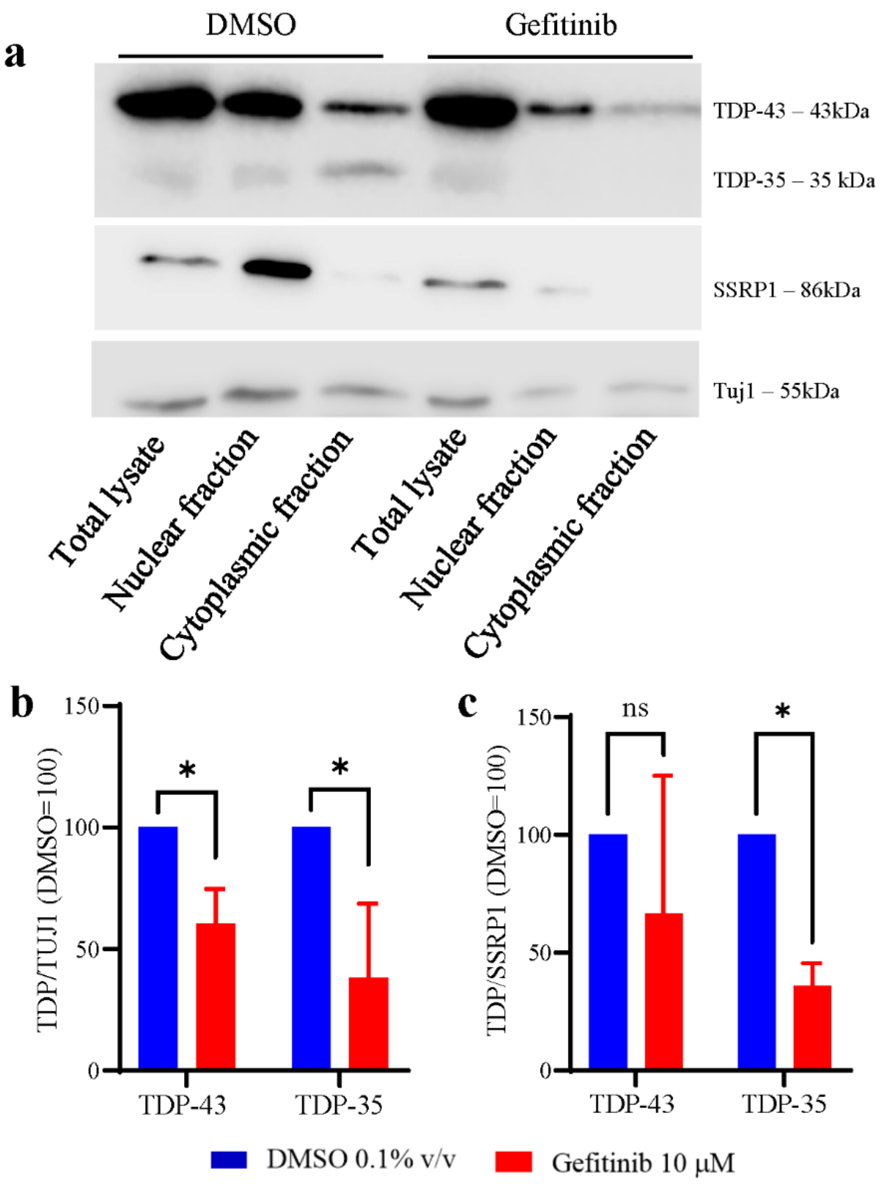


**Figure S2. Gefitinib reduces the levels of cytoplasmic TDP-35 in C9orf72-ALS iAstrocytes. A)** iAstrocytes differentiated from C9_1 were subjected to nucleocytoplasmic fractionation and the burden of TDP-43 and TDP-35 was interrogated within each fraction, alongside SSRP1 nuclear marker and Tuj1 cytoplasmic marker as loading controls. **B)** Densitometric quantification of TDP-43 and TDP-35 levels in the cytoplasmic fraction. TDP-43 and TDP-35 levels were calculated as relative to TUJ11 loading control. Data normalised to DMSO=100. Data = mean ± SD, n=4. Mann-Whitney test, *P < 0.05. **C)** Densitometric quantification of TDP-43 and TDP-35 levels in the nuclear fraction. TDP-43 and TDP-35 levels were calculated as relative to SSRP1 loading control. Data normalised to DMSO=100. Data = mean ± SD, n=4. Mann-Whitney test, *P < 0.05.

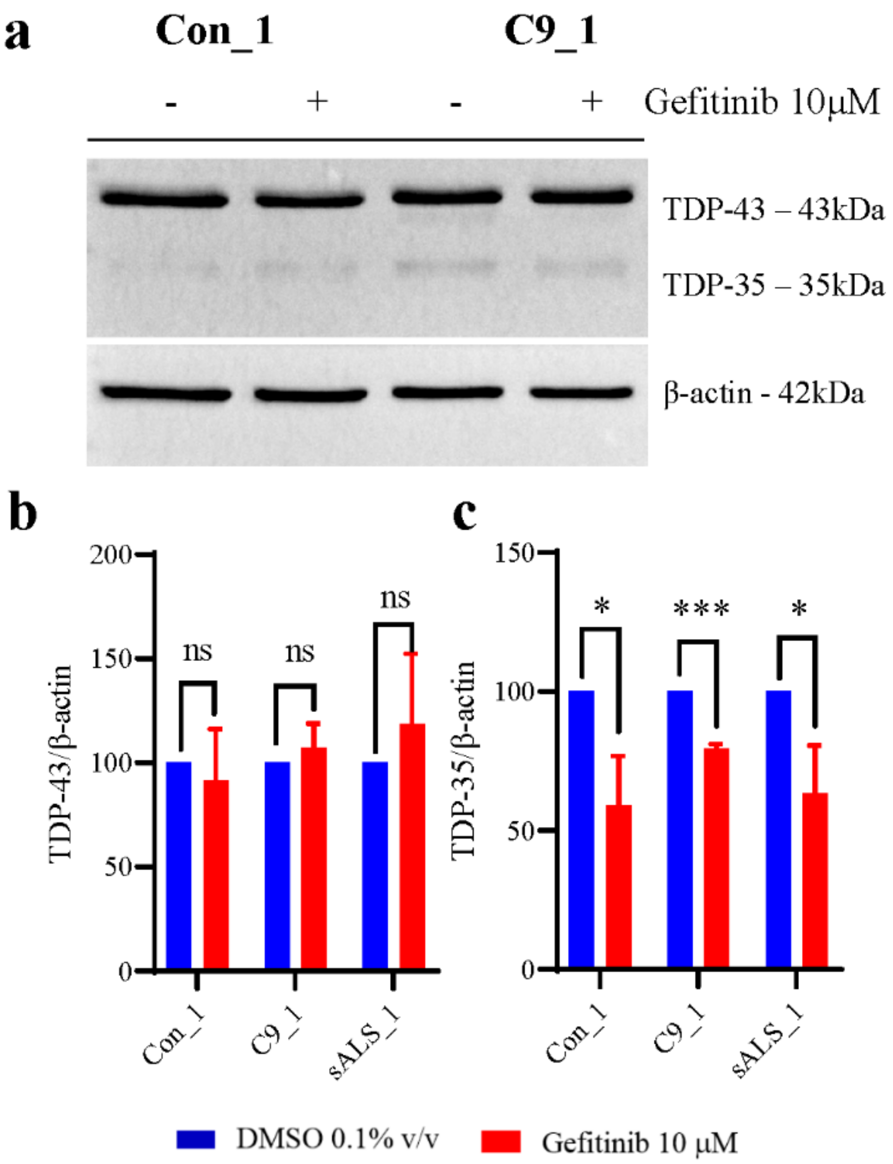


**Figure S3. Gefitinib reduces TDP-43 35 kDa fragment in C9orf72-ALS and sALS induced neurons (iNeurons). A)** Levels of TDP-43 and TDP-35 in control and C9_1 iNeurons after 48 treatment with 0.1% (v/v) DMSO and 10µM gefitinib were assessed via Western blotting. **B)** Densitometry quantification of TDP-43 and **C)** TDP-35 in iNeurons following the treatment with gefitinib. Data normalised to DMSO-treated iNeurons= 100. Data = mean ± SD, n=3. Mann-Whitney test, *P < 0.05; ***P < 0.001.

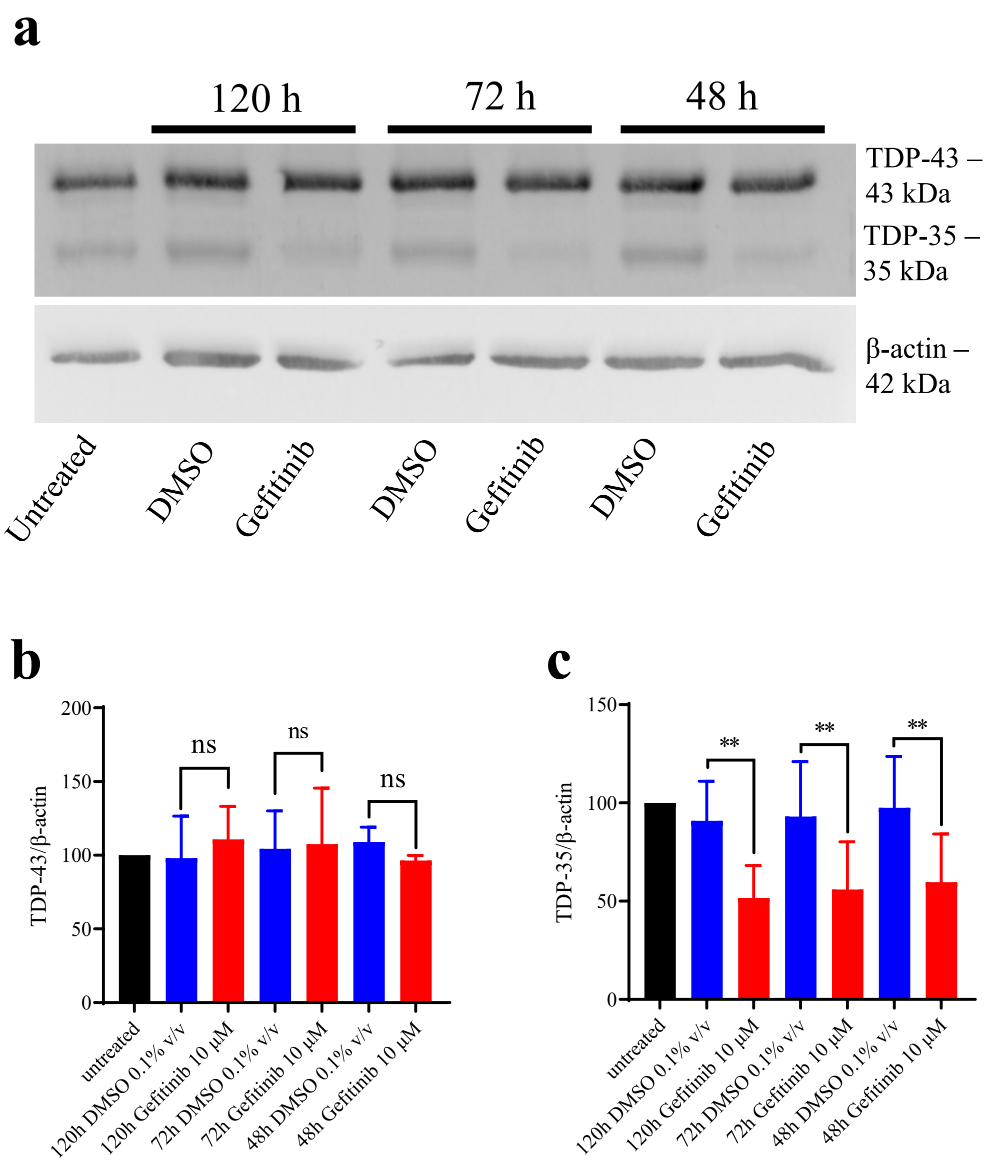


**Figure S4. Prolonged treatment with gefitinib had no further effect on TDP-43 fragmentation. A)** Representative Western blot of TDP-43 and TDP-35 expression in C9_1 iAstrocytes after 120h, 72h and 48 treatment with 0.1% (v/v) DMSO and 10µM gefitinib. **B)** Densitometry quantification of TDP-43 and **C)** TDP-35 in C9_1 iAstrocytes following the treatment with gefitinib. Data normalised to untreated iAstrocytes = 100. Data = mean ± SD, n=3. Two-way ANOVA with Tukey's multiple comparisons test, **P<0.005.

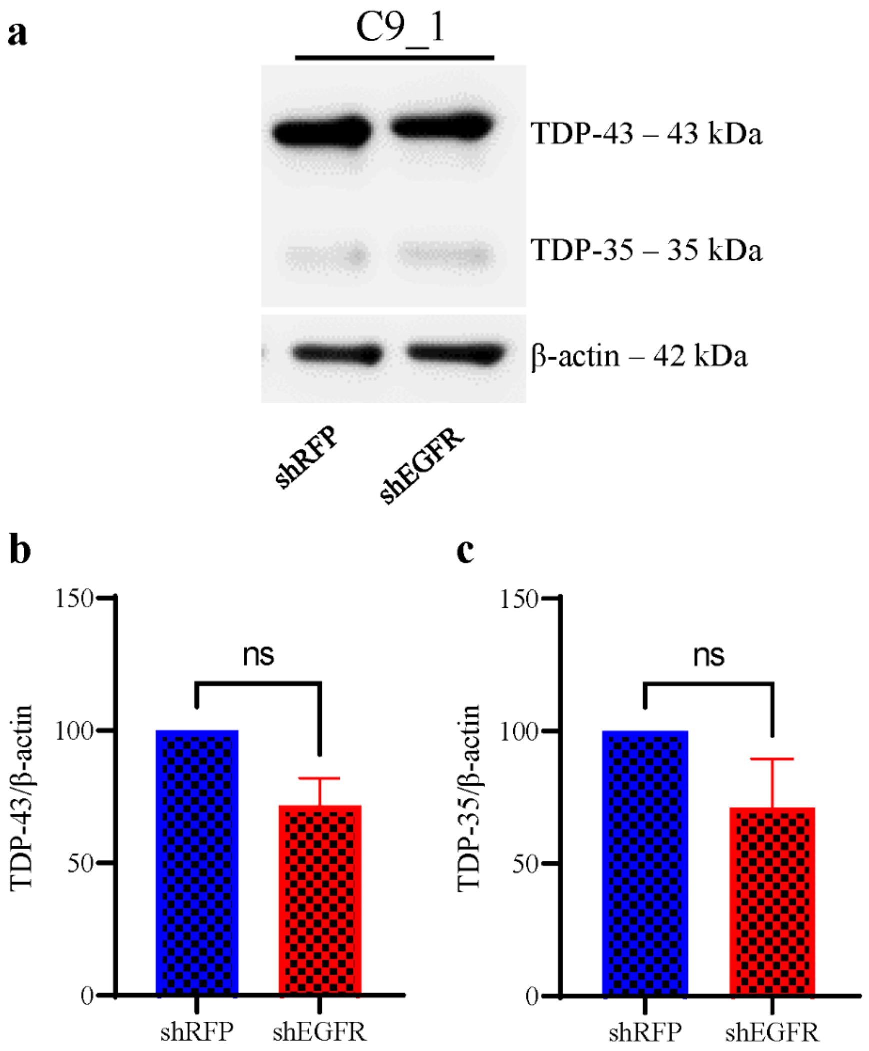


**Figure S5. Knocking-down EGFR in patient iAstrocytes has no effect on TDP-43 fragmentation. A)** Representative Western blot of C9_1 iAstrocytes treated with RFP-only vector and EGFR shRNA probed for TDP-43 and β-actin. **B)** Densitometry quantification of TDP-43 in C9_1 iAstrocytes following a treatment with shRNAs. Data = mean ± SD, n = 3. Paired t-test, ns. **C)** Densitometry quantification of TDP-35 in C9_1 iAstrocytes following a treatment with shRNAs. Data = mean ± SD, n = 3. Data normalised to RFP-treated cells = 100. Data = mean ± SD, n=3. Wilcoxon matched-pairs signed rank test, ns.

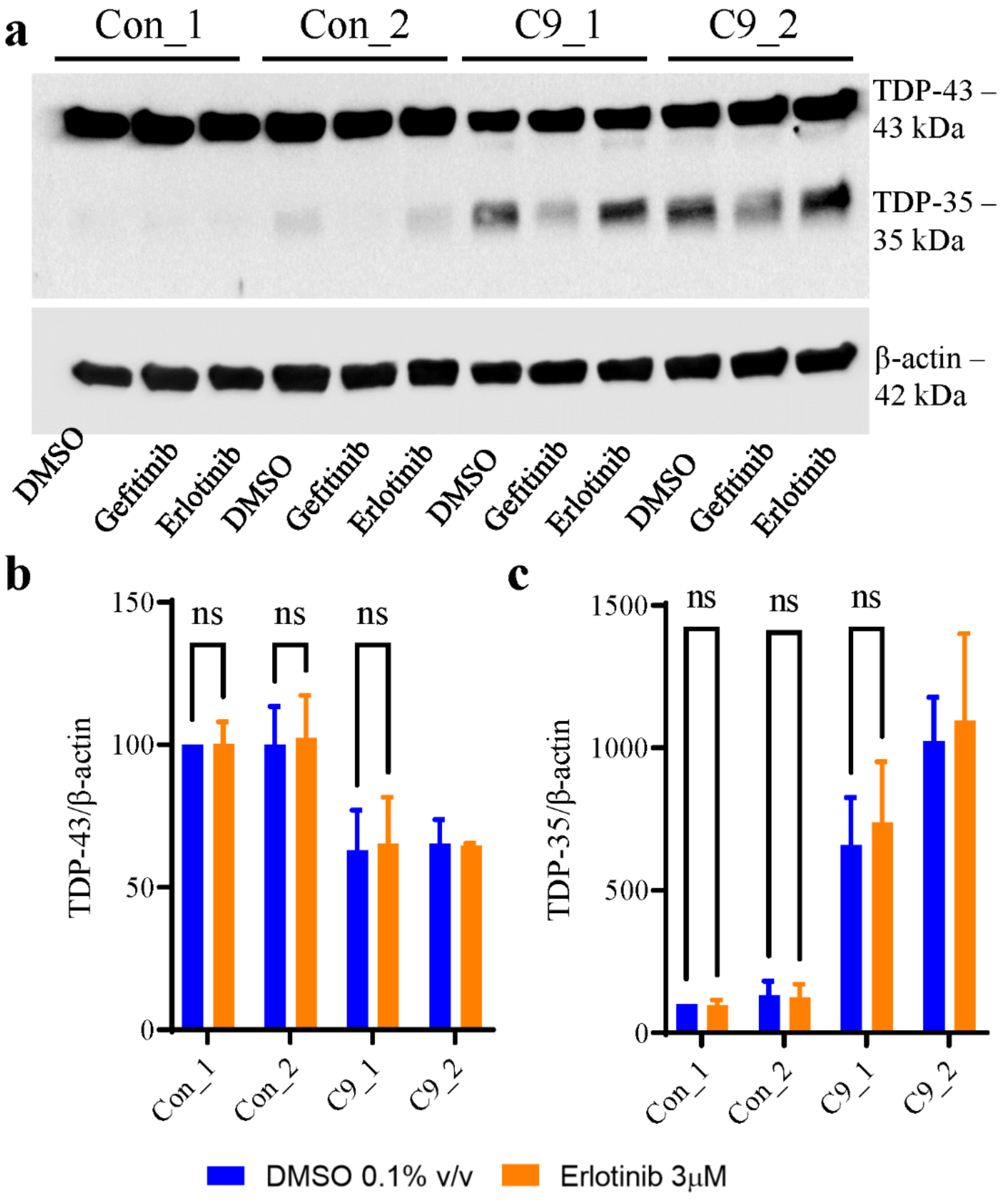


**Figure S6. Erlotinib has no effect on TDP-43 or TDP-35 fragment levels in C9orf72-ALS iAstrocytes. A)** A representative Western blot showing TDP-43 and TDP-35 in Con_1, Con_2, C9_1 and C9_2 iAstrocytes after 48 treatment with 0.1% (v/v) DMSO, 10 μM gefitinib or 3μM erlotinib. **B)** Densitometry quantification of TDP-43 and TDP-35 in iAstrocytes following the treatment with DMSO or erlotinib. Data normalised to DMSO-treated iAstrocytes of Con_1= 100. Data = mean ± SD, n=3 (except C9_2, n=2). Two-way ANOVA with Šídák's multiple comparisons test.

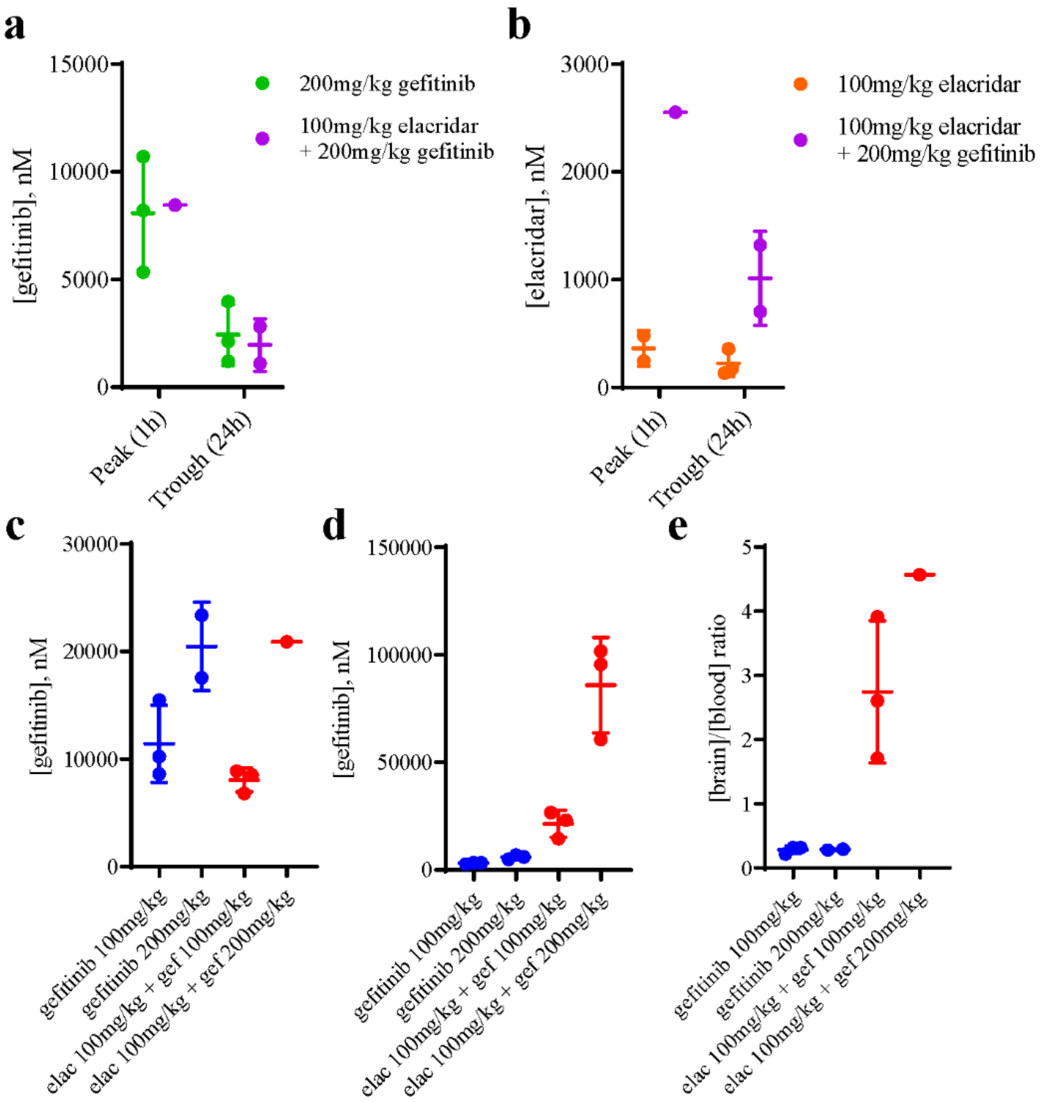


**f**

| | % free drug (*SD*) | % bound drug (*SD*) |
|---|---|---|
| Gefitinib | 0.36 (*0.01*) | 99.64 (*0.01*) |
| Warfarin | 26.43 (*2.14*) | 73.57 (*2.14*) |
| Verapamil | 2.73 (*0.19*) | 97.27 (*0.19*) |

**Figure S7. Elacridar does not affect the levels of gefitinib in blood. A)** Levels of free gefitinib (nM) at treatment peak (1h) and trough (24h) in the blood of animals dosed with 100mg/kg elacridar plus 200mg/kg gefitinib P.O., compared to animals dosed with 200mg/kg gefitinib P.O. alone. **B)** Levels of free elacridar (nM) at treatment peak (1h) and trough (24h) in the blood of animals dosed with 100mg/kg elacridar plus 200mg/kg gefitinib P.O., compared to animals dosed with 100mg/kg elacridar P.O. alone. Data = mean ± SD, n=1-3. For the pharmacokinetic analysis of gefitinib, female $SOD1^{G93A}$ C57BL/6 mice were either given a single dose elacridar (100mg/kg po, -4h), or untreated (no dose control group). Mice were then given a single oral dose of gefitinib at either 100mg/kg po or 200mg/kg po. Blood and brain were collected 2h after gefitinib administration, and gefitinib levels were measured in blood (**C**) and brain (**D**) using UHPLC - TOF mass spectrometry. (**E**) The ratio of brain/brain gefitinib concentrations was also calculated. Data = mean ± SD, n=1-3. **F)** Brain tissue protein binding was measured for gefitinib at state, by the Equilibrium Dialysis Method (EDM) followed by UHPLC - TOF mass spectrometry. The brain tissue protein binding of warfarin and verapamil were also assessed and used as medium and high binding reference standards respectively. Data = mean, n=3.

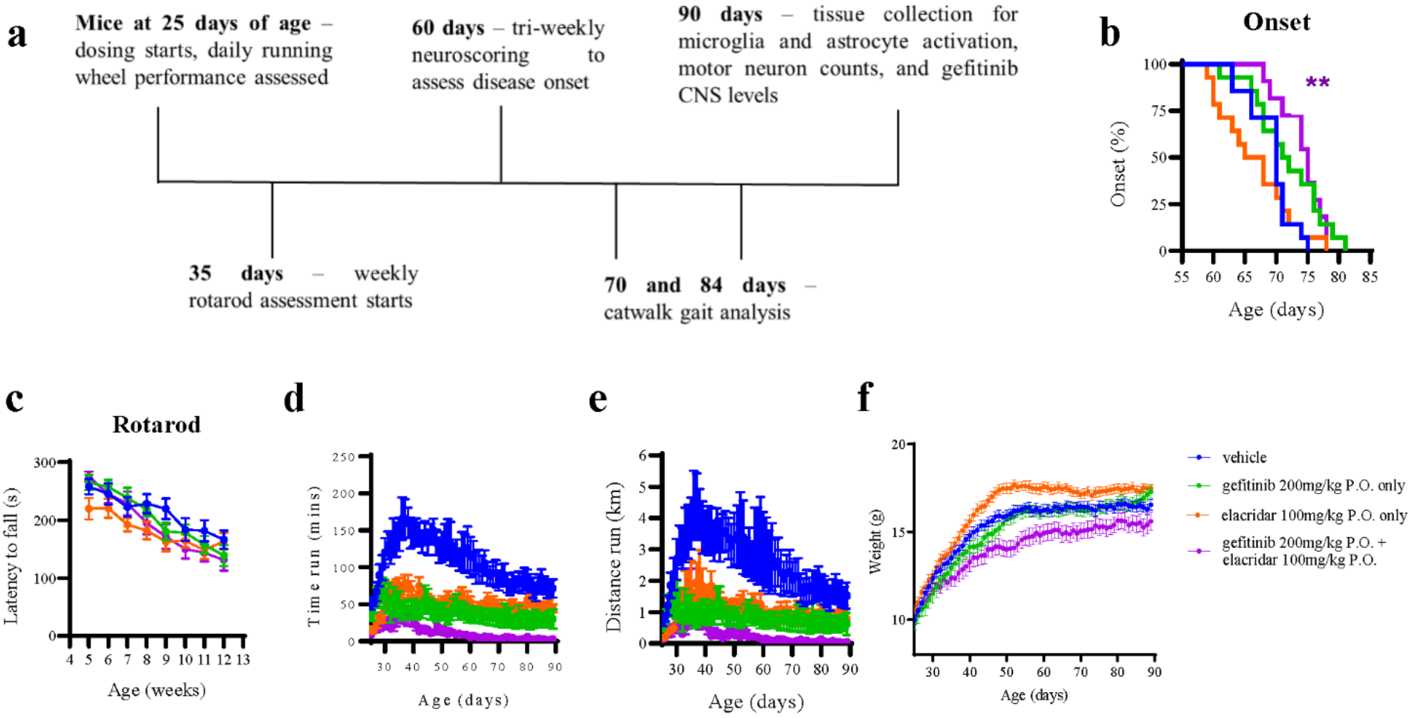


**Figure S8. High brain exposure of gefitinib delays neurological symptom onset in SOD1$^{G93A}$ mice, but does not improve motor performance – complete dataset. A)** Gefitinib *in vivo* efficacy study design. **B)** Female SOD1$^{G93A}$ mice were dosed daily from 25 days of age with vehicle control P.O. (blue), 200 mg/kg gefitinib P.O. (green), 100 mg/kg elacridar P.O. (orange), or 200 mg/kg gefitinib and 100 mg/kg elacridar P.O. (purple). Neurological symptom onset was defined when both a hind limb splay defect and hind limb tremor were observed in an animal, and the percentage of animals with onset as a function of age is plotted on a Kaplan-Meier curve. n = 11-14. One-way ANOVA with Tukey's post-hoc test; **$P < 0.01$. **C)** Rotarod performance assessed once a week from 5 weeks of age, and was measured as time to fall in seconds. Two-way ANOVA with Tukey's post-hoc test. Animals were singly housed with a Fast-trac running wheel, and time **(D)** and distance **(E)** run was recorded every 24 h. Data = mean ± SEM; n = 11-14. Repeated measures two-way ANOVA with Tukey's post-hoc test. *$P < 0.05$; **$P < 0.01$; ***$P < 0.001$; ****$P < 0.0001$. **F)** Female SOD1$^{G93A}$ C57BL/6 mice were weighted daily before dosing from 25th day of age. Animals dosed with gefitinib 200mg/kg P.O. plus elacridar 100mg/kg P.O. were significantly lighter than the vehicle-treated mice ($P<0.0001$). Data = mean ± SEM. Two-way ANOVA with Dunnett's post-hoc test. ****$P<0.0001$.

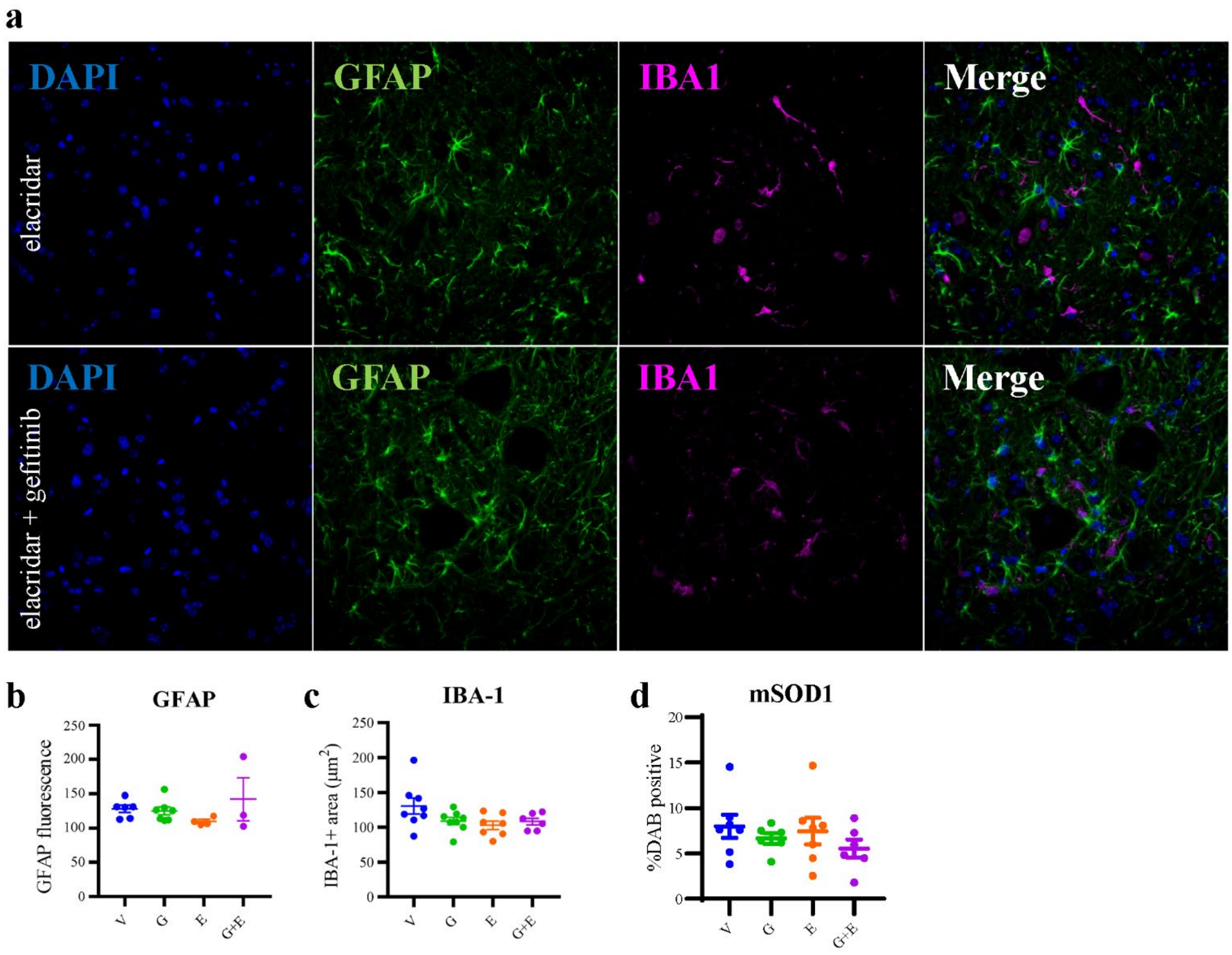


**Figure S9. High brain exposure of gefitinib reduces has no effect on glia-mediated neuroinflammation or levels of misfolded SOD1. A)** Representative images of GFAP and IBA1-stained spinal cords of $SOD1^{G93A}$ mice treated with elacridar or elacridar+gefitinib. **(B)** GFAP fluorescence intensity quantification. Data = mean ± SEM; n = 3-8. One-way ANOVA with Dunnets's post-hoc test; all comparisons ns. **C)** IBA-1-positive area quantification ($\mu m^2$). Data = mean ± SEM; n = 3-8. One-way ANOVA with Dunnets's post-hoc test; all comparisons ns. **(D)** mSOD1-positive area quantification (% DAB). Data = mean ± SEM; n = 6-7. One-way ANOVA with Dunnets's post-hoc test; all comparisons ns.
